# Structural Basis for Ca^2+^-Dependent Activation of a Plant Metacaspase

**DOI:** 10.1101/2020.03.09.983940

**Authors:** Ping Zhu, Xiao-Hong Yu, Cheng Wang, Qingfang Zhang, Wu Liu, Sean McSweeney, John Shanklin, Eric Lam, Qun Liu

## Abstract

Plants metacaspases mediate programmed cell death in development ^1,2^, biotic and abiotic stresses ^3^, damage-induced immune response ^4^, and resistance to pathogen attack ^5^. Most metacaspases require Ca^2+^ for their activation and substrate processing ^6–8^. However, the Ca^2+^-dependent activation mechanism remains elusive ^9–11^. Here we report the crystal structure of Metacaspase 4 from *Arabidopsis thaliana* (*At*MC4) that modulates Ca^2+^-dependent, damage-induced plant immune defense. The *At*MC4 structure exhibits an inhibitory conformation in which a large linker domain blocks activation and substrate access. In addition, the side chain of Lys225 in the linker domain blocks the active site by sitting directly between two catalytic residues. We show that the activation of *At*MC4 and cleavage of its physiological substrate involve multiple cleavages in the linker domain upon activation by Ca^2+^. Our analysis provides insight into the Ca^2+^-dependent activation of *At*MC4 and lays the basis for tuning its activity in response to stresses that may help engineer more sustainable crops for production of food and biofuel.

## Main

Programmed cell death is a tightly controlled process contributing to development and responses to biotic and abiotic stresses in multi-cellular organisms. In animals, caspases, upon activation by upstream signaling events, process protein substrates, leading to cell death ^12^. In plants, metacaspases were identified ^13^ to have similar roles in mediating programmed cell death in development ^1,2^, biotic and abiotic stresses ^3^, damage-induced immune response ^4^, and resistance to pathogen attack ^5^. Though caspases and metacaspases likely share common ancestors, they have evolved divergently and do not co-exist in the same organisms ^14^. Caspases are unique in metazoans while metacaspases are found in protozoans, plants, fungi, and prokaryotes.

Caspases are classified as executioners and initiators where executioner caspases are activated through proteolytic cleavage by initiator caspases ^15^. In contrast, there is no known upstream protease responsible for the activation of metacaspases. Instead, most metacaspases require Ca^2+^ for activation ^6–8^. In contrast to well-studied caspases, the activation mechanism remains elusive for *At*MC4 and other Ca^2+^-dependent metacaspases ^9–11^.

Ca^2+^ signaling controls numerous physiological activities across all kingdoms of life ^16^. In plants, Ca^2+^ flux and associated signaling events are associated with a diverse array of abiotic stresses (drought, salinity, cold, wind, wounding) and biotic stresses such as pathogen and insect attacks ^17–19^. Different stress signals encode a unique intracellular Ca^2+^ signature consisting of localization, spikes, concentration, and timing ^20^. These Ca^2+^ signatures are perceived by a diverse set of proteins that decode Ca^2+^ signals for downstream processes. In *Arabidopsis*, damage-induced intracellular Ca^2+^ flux activates Metacaspase 4 (*At*MC4) to process substrate Propep1 very precisely in time and space to initiate the immune response ^4^.

### Overall structural features

To elucidate the structural basis for Ca^2+^-dependent metacaspase activation, we determined the crystal structure of *At*MC4. *At*MC4 is a type II metacaspase, featuring a large linker domain between its p20 and p10 domains (Fig. 1a and Extended Data Fig. 1). We first solved the crystal structure of a catalytically inactive C139A mutant of *At*MC4 (Extended Data Fig. 2). The structure contains an N-terminal p20 domain, a C-terminal p10 domain, and a large linker domain that is absent in caspases, Type I and Type II metacaspases (Extended Data Fig. 1). The linker domain forms an extended patch structure consisting of a β-hairpin at its N-terminus and a large α-helical region at its C-terminus (Fig. 1a). Search of the DALI structure database server ^21^ yielded no structures similar to the linker domain. In addition to its linker domain, the p20 and p10 domains of *At*MC4 form a caspase-like core in which a two-stranded anti-parallel β-sheet from the p10 domain is in parallel with the six-stranded β-sheet from the p20 domain, sandwiched by several α-helices on both sides (Fig. 1a, b). Two catalytic residues Cys139 and His86 ^13^ are located on the p20 domain. Surprisingly, the linker domain is clamped by the two catalytic residues at the position of Lys225 (Fig. 1a), the residue critical for protease activation ^10^.

**Fig. 1.**
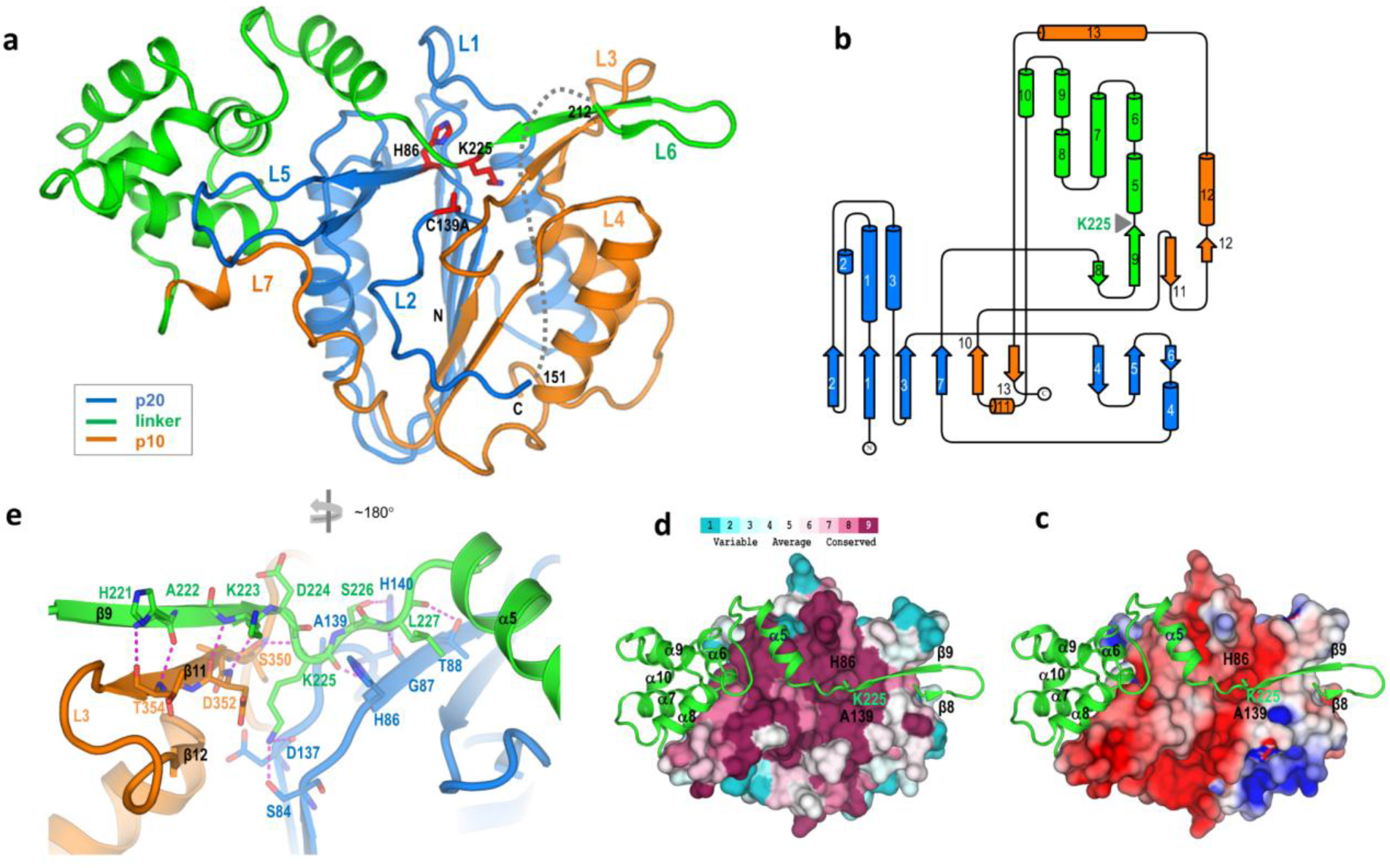
Structure of *At*MC4 and self-inhibition mechanism. **a**, *At*MC4 structure of a catalytically inactive C139A mutant. The three domains are shown as cartoons and are colored differently: p20 domain, marine; linker domain, green; p10 domain, orange. The catalytic dyad (C139A and His86) and a self-processing site (Lys225) are shown as sticks. Disordered region between residues 152 and 211 is shown as a dashed line. **b**, Topological diagram for secondary structures of *At*MC4. Coloring is the same as (**a**). **c**, Electrostatics surface of the caspase-like core. The electrostatics was calculated by using program APBS ^29^ and plotted at the level of ±4 kT/e. **d**, View of the linker domain attached on the surface of the caspase-like core with its Lys225 inserted in a conserved pocket. The conservation level is mapped to the surface: more conserved surfaces, more magenta; and more variable surfaces, more cyan. **e**, Interactions between the Lys225 region and the caspase-like core.

The caspase-like core of *At*MC4 displays structural homologies to type I Metacaspases MCA2 from *Trypanosoma brucei* ^8^ and Yca1 ^7^ from *Saccharomyces cerevisiae* but with distinctive features. The C-terminus of the p20 domain in MCA2 and Yca1 forms one and two additional β-stands, respectively, in parallel to the six-stranded β-sheet (Extended Data Fig. 3b-c). While in *At*MC4, the corresponding region is disordered, resembling human Caspase 7 (Fig. 1a, Extended Data Fig. 3a, d).

Caspases function through the activation of loops L1 through L4 to form a loop bundle structure for substrate recognition and cleavage ^15^. In human Caspase 7, the L4 loop is long; while the corresponding loop in *At*MC4 is much shorter. Instead, *At*MC4 has a long L5 loop that is very short in human Caspase 7 (Fig. 1a, Extended Data Fig. 3a, d). The L5 loop likely has a complementary role in *At*MC4 for forming a loop bundle equivalent to the L4 loop in human Caspase 7. Additionally, the L2 loop in *At*MC4 is embedded within a groove between the p20 and p10 domains (Fig. 1a); while in human Caspase 7, its L2 loop is on the surface and participates in the formation of the loop bundle (Extended Data Fig. 3d). Functional *At*MC4 is a monomer, and its L2 loop blocks the potential dimerization interface observed in Caspase 7 ^22^. Note that Ca^2+^-dependent metacaspases, including *At*MC4, MCA2, and Yca1, have a long L5 loop (Extended Data Fig. 4) and an embedded L2 loop, although the L5 loop is disordered in both MCA2 and Yca1 structures (Extended Data Fig. 3a-c).

### Self-inhibition mechanism

We used a catalytically inactive mutant (C139A) for structure determination and thus expected a structure in an inhibitory state for understanding the self-inhibition mechanism. The N-terminus of the linker domain (β8-β9-α5) crosses a highly conserved surface of the caspase-like core with residue Lys225 inserted into a conserved and negatively charged pocket (Fig. 1c, d). The pocket is comprised of conserved residues Asp137 and Ser84 from the p20 domain and Asp352 and Ser 350 from the p10 domain (Fig. 1e, and Extended Data Fig. 5a). The hydroxyl group on Ser84 and the carbonyl oxygen on Asp137 form two H-bonds with the Lys225 amide group, further stabilizing the Lys225 side chain in the pocket. These conserved residues are crucial for *At*MC4 activity; mutating any of them, except Ser350, to an alanine abolished the Ca^2+^-dependent self-cleavage (Extended Data Fig. 6).

Interactions between the linker and the caspase-like core are primarily through main-chain atoms. Indeed, residues 221-223 form an anti-parallel β-sheet with β11 and β12 associated with loop L3 (Fig. 1e). Therefore, mutating His221 and Lys223 to alanines does not affect the Ca^2+^-dependent self-cleavage (Extended Data Fig. 6c). The formation of the β-sheet structure and the locking of Lys225 in the catalytic pocket provide a structural understanding of the self-inhibition mechanism for *At*MC4. The same active site could also be the site for cleaving a substrate. Although a substrate does not necessarily have the same interactions as the inhibitory β-sheet structure, recognition through main-chain atoms suggests a low sequence specificity for becoming an *At*MC4 substrate.

### Ca^2+^-dependent self-cleavage and activation

We also sought to determine the *At*MC4 structure in an active form. However, pre-treatment of *At*MC4 with Ca^2+^ resulted in the formation of non-homogeneous fragments (Extended Data Fig. 2a) that could not be crystallized. Therefore, we first determined the wild-type *At*MC4 structure without Ca^2+^. The overall structure of the Ca^2+^-free wild-type *At*MC4 is similar to the C139A mutant but with significant conformational changes were observed for loops L2/L7 and L3/L6 (Extended Data Fig. 7).

Without Ca^2+^, the wild-type *At*MC4 structure has Lys225 locked in the active site (Fig. 2a). Cys139 and His86 have an equal distance of 3.2 Å to the carbonyl carbon of Lys225, a distance suitable for a nucleophilic attack. Nevertheless, we did not observe the bond cleavage in crystals as shown by the continuous electron densities for Lys225 (Fig. 2a). In contrast, upon treatment of wild-type *At*MC4 crystals with Ca^2+^, we observed the disappearance of electron densities for Lys225, indicating a cleavage at the position of Lys225 (Fig. 2b).

**Fig. 2.**
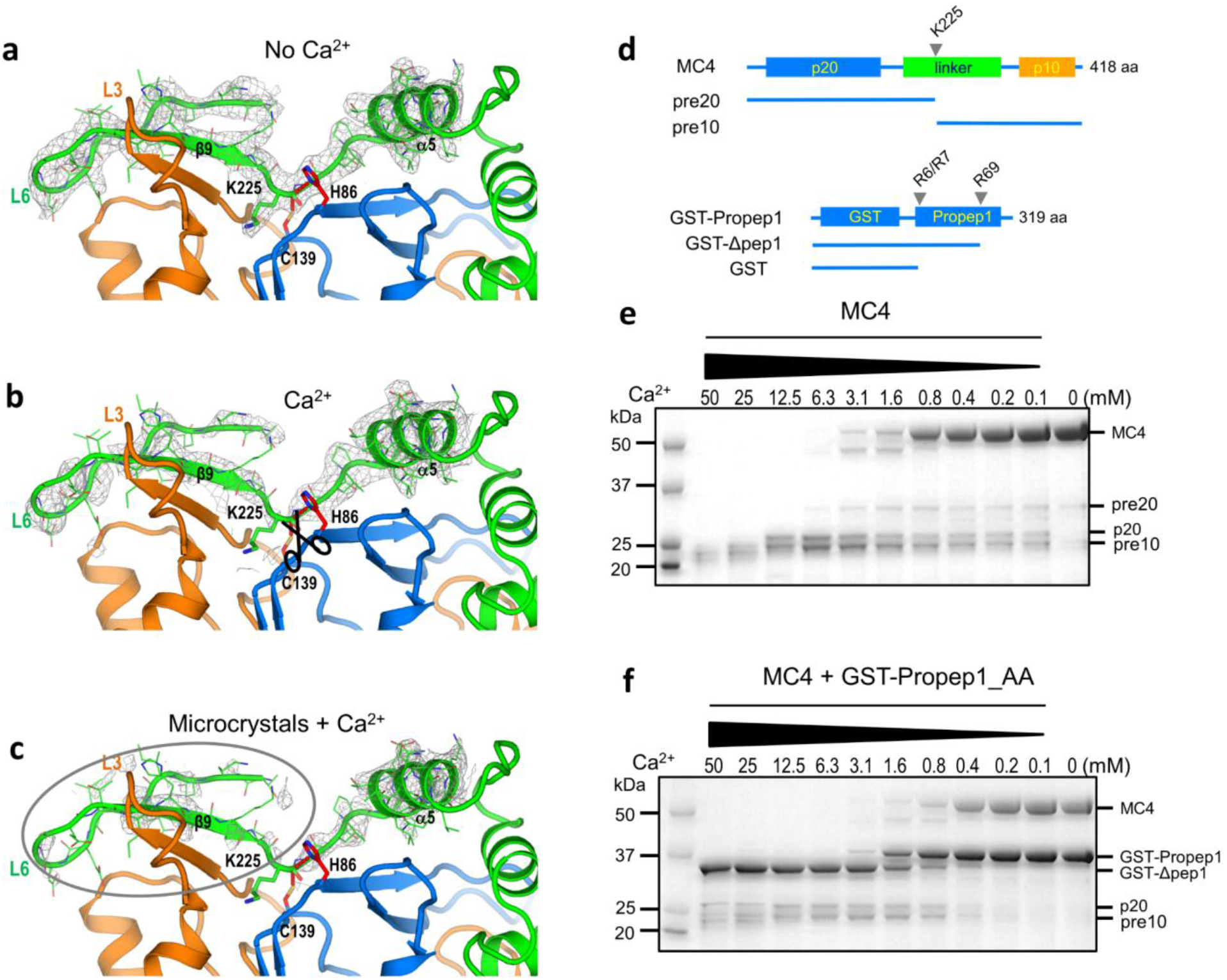
Ca^2+^-dependent self-cleavage and activation. **a-c**, Electron densities for Ca^2+^-dependent self-cleavage and activation in crystals. The electron densities were drawn as gray isomeshes contoured at 1.3σ. The distances between the catalytic dyad (His86 and Cys139) and the cleavage site of Lys225 carbonyl carbon are 3.2 Å and are shown as red dashes. **a**, Without Ca^2+^. **b**, With 10 mM Ca^2+^. **c**, With 10 mM Ca^2+^ by using microcrystals. **d**, Schematics of major fragments produced in self-cleavage of *At*MC4 and its cleavage of GST-fusion of substrate Propep1 (GST-Propep1). **e**, Ca^2+^-dependent self-cleavage and activation of *At*MC4. **f**, Ca^2+^-dependent processing of GST-Propep1 by *At*MC4. Here a R6A/R7A double mutant in Propep1 is used to highlight the Ca^2+^-dependent cleavage at the site of Arg69. Cleavage of the wild-type GST-Propep1 is shown in Extended Data Fig. 8e where higher concentration of Ca^2+^ cleaves Propep1 at an additional site of R6/R7.

With a prolonged Ca^2+^ treatment or with higher Ca^2+^ concentrations, the crystals deteriorated and lost diffraction perhaps due to large conformational changes after the Lys225 cleavage. We hypothesized that microcrystals may allow for large conformational changes without compromising their diffraction. We therefore prepared wild-type *At*MC4 microcrystals and solved Ca^2+^-treated structure by using the method we recently developed ^23^. Ca^2+^-treated microcrystals changed the space group, shrunk the unit cell dimensions, and diffracted X-rays better than the non-treated large crystals (Extended Data Table 1). In the solved Ca^2+^-treated structure, we observed the disappearance of electron densities for the entire N-terminus of the linker domain (linker-N), including Lys225, β9, and loop L6 (Fig. 2c). Therefore, after the first cleavage at Lys225, there appears to be a large conformational change at the N-terminus of the linker domain. We imagine that in solution, the effects from Ca^2+^ treatment will trigger significant conformational changes, which could facilitate the cleavage of additional sites (Arg180, Arg190, and Lys210) in the N-terminus of the linker domain ^4^ on the pathway toward full protease activation.

To understand additional steps required for *At*MC4 activation, we performed comparative Ca^2+^-dependent cleavage of its physiological substrate Propep1 ^4^. For simplicity, we used a GST-fusion of Propep1 (GST-Propep1) (Fig. 2d, Extended Data Fig. 5b) for testing its cleavage by *At*MC4. Self-cleavage of *At*MC4 is Ca^2+^-concentration dependent. Higher concentrations of Ca^2+^ produce smaller fragments (Fig. 2e). *At*MC4 cleaves GST-Propep1 also in a Ca^2+^-concentration dependent manner (Fig. 2f and Extended Data Fig. 8e). Ca^2+^ concentrations of 0.4-0.8 mM can initiate the cleavage at position Arg69 to produce Pep1 peptides that can activate a plant immune response by forming a complex with its receptor PEPR1 ^4,24^. At a Ca^2+^ concentration of 12.5 mM or higher, GST-Δpep1 (Fig. 2d) is further processed at the position of Arg6 or Arg7 (Extended Data Fig. 8e). A GST-Propep1 R6A/R7A double mutant prevented this further cleavage at higher Ca^2+^ concentrations (Fig. 2f).

### Loop L5 is critical for Ca^2+^-dependent activation and substrate cleavage

In the *At*MC4 structure, the L5 loop of the p20 domain lies in a positively charged concave surface mainly formed by the linker domain (Fig. 3a). Sequence alignments for the L5 loop region for nine *Arabidopsis* metacaspases (*At*MC1-9) revealed a cluster of negatively charged residues in *At*MC4 to *At*MC8, but not in *At*MC9 whose activation does not require Ca^2+^ (Fig. 3b) ^9^. In the *At*MC4 structure, the 96EDDD99 segment has three H-bond interactions with Lys276, Lys320, and Ala325 in the linker domain (Fig. 3c). We suggest that these negatively charged residues are involved in Ca^2+^-dependent protease activation. To test this hypothesis, we made a tetra-mutant by mutating each of them to an alanine. With increased Ca^2+^ concentration, the tetra-mutant can be partially self-processed to a major fragment of pre10 (produced after cleavage at the Lys225 position) (Fig. 2d, Extended Data Fig. 8a), but lacks further cleavage at a Ca^2+^ concentration of 12.5 mM or higher (Fig. 3d). This tetra-mutant could not cleave the GST-Propep1 effectively even at a Ca^2+^ concentration of 25 mM or higher (Fig. 3d).

**Fig. 3.**
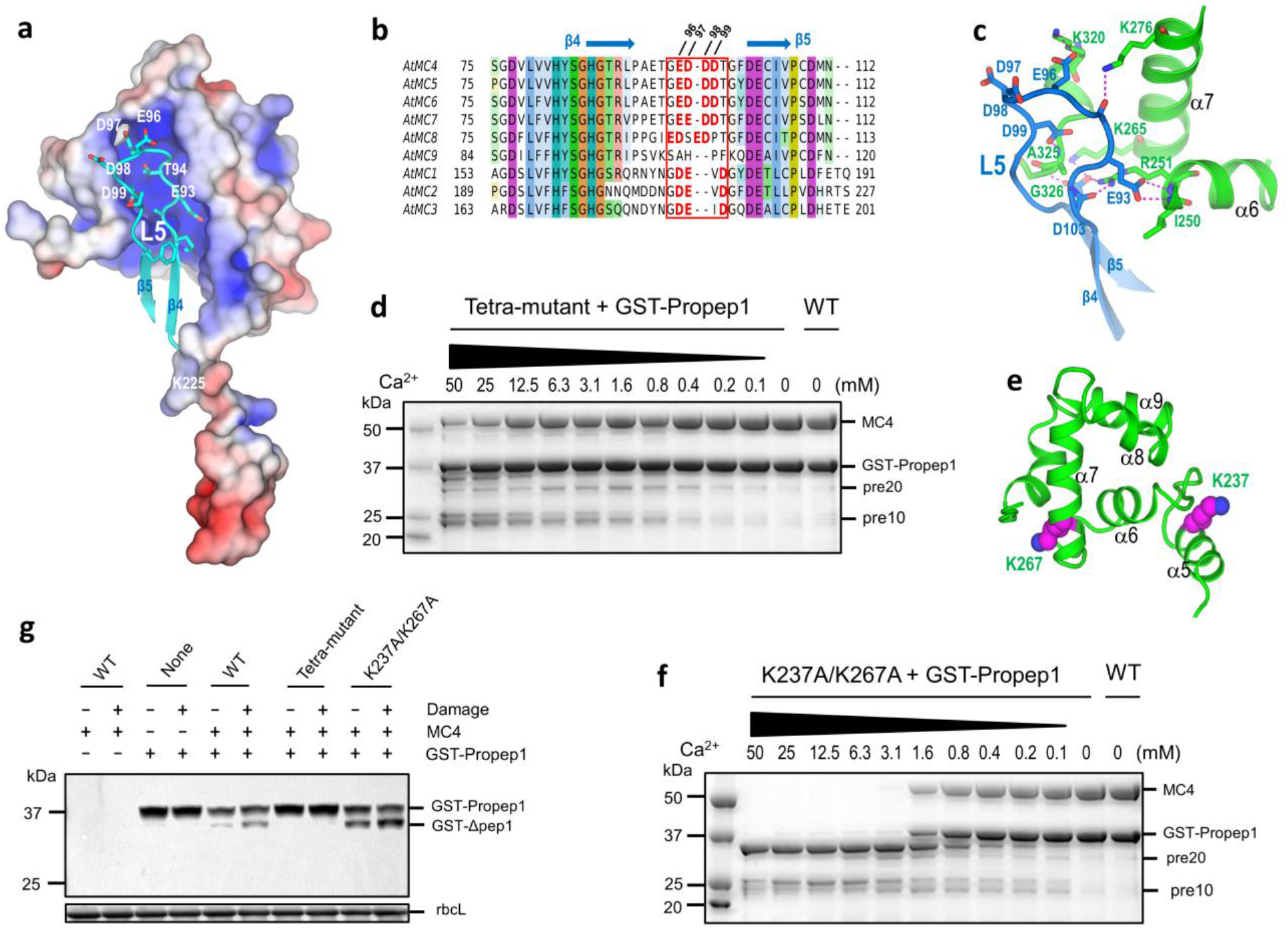
The roles of L5 loop and two self-cleave sites in the linker-C domain in Ca^2+^-dependent *At*MC4 activation and substrate processing. **a**, Loop L5 (cyan) is imbedded in a positively charged pocket defined by the linker domain. The electrostatics were calculated by APBS and plotted at the level of ± 4 kT/e. **b**, Sequence alignment of nine *Arabidopsis* metacaspases (*At*MCs) for the L5 loop region. Highlighted residues (bold in red) are negatively charged residues present in all *At*MCs except *At*MC9 whose activation is Ca^2+^ independent. **c**, Interactions of the L5 loop with the linker domain. H-bonds are shown as dashed lines and colored in magenta. **d**, A tetra-mutant in the L5 loop is deficient to cleave its substrate GST-Propep1 *in vitro*. **e**, Location of two self-cleavage sites (Lys237 and Lys267) in the α-helical region of the linker domain (Linker-C region). **f,** A K237A/K267A double mutant cleaves GST-Propep1 in a less Ca^2+^-dependence *in vitro*. **g**, *In vivo* damage-induced GST-Propep1 processing by wild-type *At*MC4 and its mutants in transfected tobacco leaves. Western blot used anti-GST antiserum. For protein loading control, the blot is stained by Ponceau Red and the band for rbcL is indicated.

In the Ca^2+^-treated structure, we did not observe a Ca^2+^-binding site. This is not surprising as the Ca^2+^ binding affinity to *At*MC4 is likely to be low ^10^. To understand the structural basis for the role of Ca^2+^ in *At*MC4 activation, we aligned the *At*MC4 structure with the MCA2 structure in which a Sm^3+^ is coordinated with four aspartate residues and two water molecules ^8^. However, in the *At*MC4 structure loops L2 and L7 appear to sterically clash with the two water molecules and Arg120 (Asp220 in MCA2) does not favor a coordination with Sm^3+^ (Extended Data Fig. 3e). To test whether Sm^3+^ can bind to this site in *At*MC4, we soaked wild-type *At*MC4 crystals with Sm^3+^ and solved the structure. In the solved structure, we did not observe a Sm^3+^-binding site as observed in the MCA2 structure. Instead, we found that Sm^3+^ binds to Asp98 in L5 loop together with Asp74 from a symmetry-related molecule (Extended Data Fig. 3f).

Sm^3+^ is a Ca^2+^ surrogate; and its affinity to Asp98 might provide a clue for understanding the role of Ca^2+^ in *At*MC4 activation and substrate processing. We thus made a D98A mutant and tested its Ca^2+^-dependent self-cleavage and cleavage of GST-Propep1. The self-cleavage in D98A is Ca^2+^-dependent until the production of the p20 and pre10 fragments (Extended Data Fig. 3g). D98A remains active to process substrate GST-Propep1 (Extended Data Fig. 3h). However, compared to the wild-type *At*MC4 (Extended Data Fig. 8e), D98A is much less active in cleaving substrate. Likely, interactions between Ca^2+^ and the negatively charged L5 loop may destabilize the electrostatic interactions between the L5 and the linker domain (Fig. 3a), thereby promoting the displacement of the linker domain from the caspase-like core toward *At*MC4 activation. It is also possible that Ca^2+^ might mediate interactions between the L5 loop of *At*MC4 and its substrate as implied by the Sm^3+^-mediated interactions (Extended Data Fig. 3f). Nevertheless, the tetra-mutant is deficient in further self-processing or cleaving its substrate; and the L5 loop thus likely has a critical role in *At*MC4 activation and effective substrate cleavage.

### Mechanism of Ca^2+^-dependent multiple cleavages and activation

To identify these self-cleavage sites in the linker domain during Ca^2+^-dependent *At*MC4 activation (Fig. 2e), we dissolved wild-type *At*MC4 crystals in a buffer containing 0.2 mM Ca^2+^ and used mass spectrometry to identify possible cleavage sites. In addition to the previously identified sites of K225, R180, R190, and K210 in the linker-N region ^4^, we identified Lys237 and Lys267 in two α-helices of the linker-C region (Fig. 3e, Extended Data Table 2). Although mutating each of them to an alanine did not change the self-cleavage activity significantly (Extended Data Fig. 8b-c), the K237A/K267A double mutant strongly reduced the Ca^2+^-dependent self-cleavage at the p20 and pre10 stage (Extended Data Fig. 8d). Interestingly, this double mutant appears to be more responsive to Ca^2+^ in the cleavage of GST-Propep1. Even with a low concentration of 0.1 mM Ca^2+^, GST-Propep1 processing can be observed (Fig. 3f). We attribute this enhanced sensitivity to either reduced electrostatics interactions between loop L5 and the linker domain or increased flexibility of the linker-C region in the double mutant.

Our structural and functional analyses of *At*MC4 lead us to propose a Ca^2+^ dependent, multi-cleavage process for metacaspase activation as illustrated in Fig. 4. Under resting conditions, the linker domain blocks the active site and the loops L3 and L5, maintaining the zymogen in an inactive state (Fig. 4a). Starting with the inactive zymogen, a Ca^2+^ concentration at sub-millimolar levels can initiate cleavage at Lys225 (Figs. 4b, 2e), leading to increased disorder of the linker-N region (Fig. 2c) and cleavage at additional sites such as R180, R190 and K210 in the linker-N ^4^, producing mainly p20 and pre10 (Figs. 4c, 2e). After the release of the linker-N region, the active site of *At*MC4 is available to process substrates such as Propep1 (Fig. 4d) to produce the Pep1 elicitor and trigger the immune response downstream ^4^. In addition, the release of the linker-N region will destabilize the β11-β12 hairpin (Fig. 1e), forming a long and disordered L3 loop. Notably the analogous L3 loop is disordered in the MCA2 and Yca1 structures that do not have an inhibitory linker domain (Extended Data Figs. 3b-c). Further cleavage at positions of Lys237 and Lys267 involves the unfolding of α helices in the linker-C region (Figs. 3e, 4e), which likely destabilizes the L5 loop that would undergo conformational changes to form a proposed loop bundle with loop L3 for a fully activated enzyme (Fig. 4f). The loop bundle might be important in substrate processing through an induced-fit mechanism as observed in the inhibitory structure (Fig. 1a) and as proposed for caspases ^25^.

**Fig. 4.**
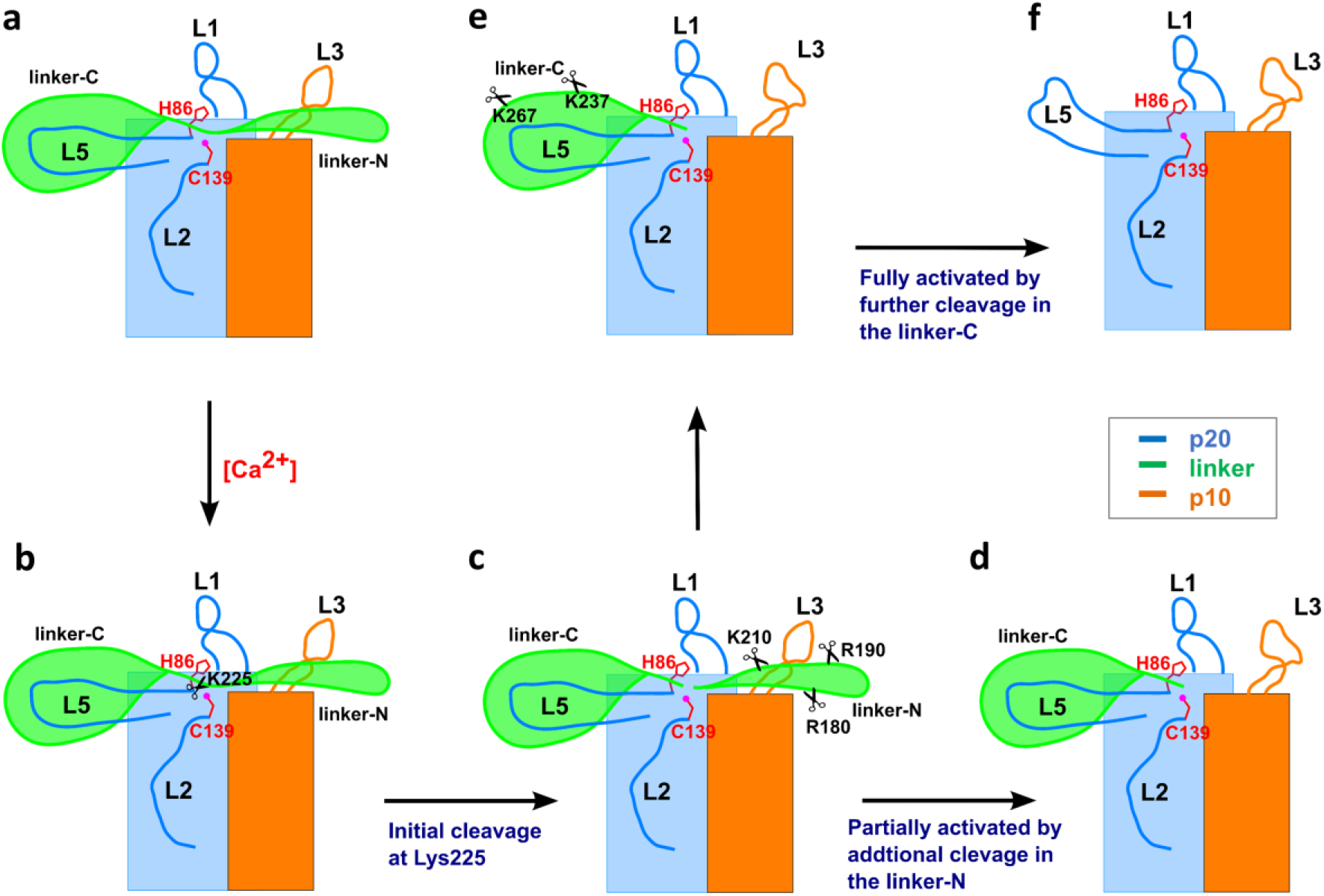
Proposed mechanism of Ca^2+^-dependent *At*MC4 activation. Two rectangular boxes are for p20 (blue) and p10 (orange) domains. The linker domain is consisted of linker-N (N-term) and Linker-C (C-term) separated by residue Lys225. Two catalytic residues Cys139 and His86 are shown as red sticks. Cleavage sites Lys237 and Lys267 were identified in the study; cleavage sites Arg180, Arg190, and Lys210 were from Hander, et al. ^4^ and were confirmed in this study. **a**, Inactive form. **b-c**, Initial cleavage at Lys225. **d**, Additional cleavage in the linker-N for partial activation. **e**, Further cleavage in the linker-C for full activation.

### Damage-induced substrate cleavage *in vivo*

To validate our structural and *in vitro* analyses in damage-induced immune-response *in vivo*, we transiently expressed wild-type *At*MC4 or its mutants, with GST-*At*Propep1 in tobacco leaves by agroinfiltration and examined their efficacy in the cleavage of GST-*At*Propep1. With agrobacterium infiltration, both wild-type *At*MC4 and K237A/K267A can cleave GST-*At*Propep1 to produce Pep1 peptides; while increased cleavage was seen for the K237A/K267A mutant *in vivo* (Fig. 3g, Extended Data Fig. 8f). This is consistent with the reduced Ca^2+^-dependence observed *in vitro* for substrate processing (Fig. 3f). The observation of increased substrate-cleaving activity by the K237A/K267A mutant suggests a negative regulation of substrate cleavage by these two sites. Damage further enhanced the cleavage of GST-*At*Propep1, likely through elevated Ca^2+^ concentration ^4^. In contrast, the tetra-mutant cannot effectively process GST-*At*Propep1 (Fig. 3g), consistent with *in vitro* observations described above.

## Discussion

Upon exposure to adverse environments such as salinity, drought, cold, wind, physical damage, and pathogen attack, proper responses to these stresses are traits that will be critical for breeding sustainable crops. Diverse types of plant stresses have been associated with tightly controlled Ca^2+^ flux and an elevation of Ca^2+^ concentration in the cytosol ^17^. *At*MC4 is localized in cytosol where Ca^2+^ concentration is strictly maintained at about 100 nM under resting conditions. Therefore, it is likely that *At*MC4, though constitutively and abundantly expressed in *Arabidopsis*, is kept in a resting state as shown in our Ca^2+^-free structures. In plants, there are multiple Ca^2+^ stores (vacuole, ER, Golgi and cell wall) where Ca^2+^ concentration varies between sub-mM to mM range ^26^. Under physical damage, pathogen attack or other stresses, Ca^2+^ fluxes from these stores can produce different local Ca^2+^ concentrations to activate metacaspases such as *At*MC4 to initiate different extents of self-cleavage for the appropriate immune responses ^4,27,28^. We thus propose that metacaspases function as a Ca^2+^-signature decoder to transduce Ca^2+^ signals to activate distinct response pathways. Understanding the structural basis for Ca^2+^-dependent metacaspase activation may enable the fine tuning of its activity in response to abiotic and biotic stresses for engineering more sustainable crops for production of food and biofuel.

## Acknowledgments

We thank Wayne A. Hendrickson for suggestions, Jianqiao Fortin for help with *At*MC4 constructs, John Haley for help with mass spectrometry, and staff at the NSLS-II beamlines FMX and AMX for their assistance in data collection. This work was supported by the U.S. Department of Energy (DOE), Office of Biological and Environmental Research. Structure determination used a method that was supported in part by NIH grant GM107462. E. L. was supported by NSF grant IOS-1258071. The work used National Synchrotron Light source II (NSLS-II) which is supported in part by the U.S. DOE Office of Basic Energy Sciences under contract number DE-SC0012704. Beamlines FMX and AMX are supported by NIH P41GM111244 and by the DOE Office of Biological and Environmental Research.

## Author contributions

P.Z., X.Y., J.S., E.L., and Q.L. designed the study and experiments. P.Z., X.Y., C.W., Q.Z, W.L. and Q.L. performed the experiments. P.Z., X.Y., S.M. J. S., E.L, and Q.L. analyzed the data. E.L. and Q.L. wrote the manuscript with help from the others.

## Additional information

**Supplementary information** is available for this paper at https://xxxxx

**Correspondence and requests for materials** should be addressed to J.S., E.L. or Q.L.

## Data availability

Atomic coordinates and structure factor files have been deposited in the RCSB Protein Data Bank (PDB) under the accession code XXXX for the C139A mutant, XXXX for the wild-type, and XXXX for the Ca^2+^-treated wild-type microcrystals.

## Methods

### Protein production for *At*MC4

The full-length, wild-type *AtMC4* (residues 1-418) was subcloned from clone 183F14 (Accession number: H37084) into the BamHI and XhoI sites of pET23a vector (Novagen) using standard PCR-based protocols. Mutagenesis of *At*MC4 was performed using a one-step PCR method ^30^.

Proteins were overexpressed in *Escherichia coli* BL21 (DE3) pLysS at 22 °C for 3-6 hrs induced by addition of 0.4 mM IPTG (final) to the cell culture with an A600 of 0.4–0.6. Harvested cells were resuspended in extraction buffer that contains 25 mM Tris, pH 7.6, 250 mM NaCl, 0.5 mM TCEP, 5% glycerol and protease inhibitors. Cells were lyzed by using an EmulsiFlex-C3 Homogenizer (Avestin, Ottawa, Canada). After centrifugation at 18,000xg for 1 hr, the supernatants were collected for a three-step purification by nickel-nitrilotriacetic acid affinity chromatography (HisTrap FF column, GE Healthcare, Inc.), ion exchange chromatography (HiTrap Q HP column, GE Healthcare, Inc.), and gel filtration (Superdex-200 10/300 GL column, GE Healthcare, Inc). Purified proteins were concentrated by using an Amicon Ultra-15 centrifugal filter (Milipore, Inc).

### Protein production for GST-Propep1

The coding sequence for the full-length AtPropep1 (residues 1-92) (clone 06-11-N09 from RIKEN, Japan) was subcloned into the BamHI and XhoI sites of pGEX-4T-1 vector (GE Healthcare, Inc.) using standard PCR-based protocols. The protein was overexpressed in *Escherichia coli* BL21 (DE3) pLysS at 16 °C for 20 hrs induced by addition of 0.2 mM IPTG (final) to the cell culture with an A600 of 0.4–0.6. Harvested cells were resuspended in extraction buffer that contains 25 mM Tris, pH 7.6, 150 mM NaCl, 10 mM DTT, 5% glycerol and protease inhibitors. After cells were lyzed by using an EmulsiFlex-C3 Homogenizer (Avestin, Ottawa, Canada), Triton X-100 (Sigma) was added to lysates to a final concentration of 1% and stirred at 4 °C for 1 hr. After centrifugation at 18,000xg for 1 hr, the supernatant was purified by using Glutathione Sepharose 4B resin and PD10 column according to the manufacturer’s protocols (GE Healthcare, Inc.). The resins were washed with wash buffer (25 mM HEPES, pH 7.6, 150 mM NaCl, 10 mM DTT, 0.2 mM AEBSF, and 5% Glycerol). GST-Propep1 protein was eluted by wash buffer supplemented with 10 mM glutathione and 0.1% Triton X-100.

### Crystallization

Crystallizations of C139A and wild-type *At*MC4 were performed by using the vapor diffusion hanging drop method. For crystallization of the C139A mutant and its SeMet substitution of *At*MC4, 1 μL of 30 mg/ml protein was mixed with an equal volume of precipitant that contains 100 mM sodium acetate, pH 4.6 and 2.1 M ammonium sulfate. For crystallization of wild-type *At*MC4, 1 μL of protein (20 mg/ml) was mixed with an equal volume of precipitant (100 mM sodium cacodylate, pH 6.4, 2.1 M ammonium sulfate). For cryo-crystallography, 10% glycerol was supplemented to the precipitates to form cryoprotectants. Crystals were transferred into their respective cryoprotectants prior to be cryocooled into liquid nitrogen for cryogenic data collection.

### Crystal soaking experiments

Freshly grown wild-type *At*MC4 crystals were harvested in cold room. The Ca^2+^ soaking solution contains 100 mM sodium cacodylate, pH 6.4, 2.1 M ammonium sulfate, 0.2, 1, or 10 mM CaCl_2_, and 10% glycerol. After addition of crystals into the soaking solution, we sealed the soaking drops and moved them to room temperature for 10 min. We then moved crystals back to cold room for freezing them into liquid nitrogen. The soaking of wild-type *At*MC4 crystals by SmCl_3_ was performed the same as we did for the Ca^2+^ soaking.

To see conformational changes after the initiation of the Lys225 cleavage, we utilized microcrystals for treatment by 10 mM Ca^2+^. We transferred large crystals into small drops of the soaking solution. We then smashed crystals into microcrystal pieces, sealed the soaking drops and moved them to room temperature for 10 min. We then moved microcrystals back to cold room. To manipulate microcrystals for microdiffraction data collection, we used a pipette to aspire microcrystal slurries, put them on the custom-made MiTiGen wellmounts ^23^, and flash-frozen them into liquid nitrogen.

### Diffraction data collection and analysis

Diffraction data were collected at NSLS-II beamline FMX with an Eiger 16M detector and beamline AMX with an Eiger 9M detector, both under a cryogenic temperature of 100 K. To collect Ca^2+^-treated microcrystal data sets on wellmounts, we used raster scans with a step size of 5 μm to find positions with diffracting crystals, and selected these positions for collection of 20° of rotation data from each position. All data sets were indexed and integrated by DIALS ^31^ and scaled and merged by CCP4 program AIMLESS ^32^. Data collection and reduction statistics for single- and multi-crystal data sets are listed in Extended Data Table 1.

### Structure Determination

The C139A mutant structure was determined by single-isomorphous replacement with anomalous scattering (SIRAS). To enhance anomalous signals from Se sites at a low resolution of 4 Å, we used an iterative crystal and frame rejection technique that we developed for microcrystals ^23^. The assembled data were used for substructure determination by program SHELXD ^33^. Se-substructures were used for phasing in SHARP/autoSHARP ^34^ by single isomorphous replacement with anomalous signals. Initial model was built automatically by program BUCCANEER ^35^. Further model building and refinements were respectively performed in phenix.refine ^36^ and COOT ^37^. There are two *At*MC4 molecules in a.u. We used non-crystallographic symmetry (NCS) for restraints and TLS parameters were used to model anisotropy.

In addition to the C139A mutant, we also crystallized wild-type *At*MC4. These crystals appeared within a couple of days; but they then underwent an aging process and lost diffraction with a prolonged growth after three days. Therefore, only fresh wild-type crystals were used for structure analysis. The wild-type *At*MC4 structure was determined by molecular replacement with the C139A structure as a start model. Structures were rebuilt and refined iteratively in COOT and phenix.refine, respectively. NCS restraints were used to improve stereochemistry. The stereochemistry of refined structures was validated with PROCHECK ^38^and MOLPROBITY ^39^ for quality assurance. Data statistics for refinements were listed in Extended Data Table 1.

### Structure determination from Ca^2+^-treated microcrystals

We solved the Ca^2+^-treated structure by combining partial data from 12 microcrystals using a modified data assembly method that we have developed ^23^. Briefly, we collected 132 partial data sets, each from a Ca^2+^-treated microcrystals. Based on unit cell variation analysis ^40^, we classified them into 20 groups to reject data sets with large unit cell variations. For each individual group, we scaled and merged their group members by using CCP4 program AIMLESS. Most of the merged data sets are incomplete. We thus selected 4 groups (11, 12, 13 and 15) with a completeness greater than 90% for structure determination by molecular replacement followed by model building and refinement. Among these merged data sets, group 11 displays the lowest refined R free (0.35) and contains data merged from 14 microcrystals. This data was further optimized by using the iterative crystal and frame reject method ^23^. The optimized data used 338 frames from 12 microcrystals. This data was used for further structural refinement with a final refined R free of 0.32. The data collection and refinement statistics for the 12-microcrystal data were listed in Extended Data Table 1.

### *In vitro* cleavage assays

*At*MC4 self-cleavage activity was measured by incubation of 5 uM of the purified *At*MC4 or its mutants for 10 min at room temperature in 25 uL reaction solution containing 25 mM HEPES, pH 7.6, 250 mM NaCl, 0–50 mM CaCl_2_, and 0.5 mM TCEP (tris 2-carboxyethyl phosphine). Cleavage of substrate GST-Propep1 by *At*MC4 and its mutants were measured by incubation of 5 uM of purified *At*MC4 and 10 uM of purified GST-Propep1 for 30 min at room temperature in 25 uL reaction buffer containing 25 mM HEPES, pH 7.6, 250 mM NaCl, 50 mM CaCl_2_, and 0.5 mM TCEP. Reactions were stopped by the addition of SDS-PAGE sample buffer supplemented with 50 mM EDTA. Proteins were separated in 4-20% gradient precast PAGE gels (Genscript) and stained by Coomassie blue.

### *In vivo* damage assays

*At*MC4 and its mutants were amplified from their corresponding expression vectors and cloned into pCR8/GW-TOPO to generate donor vectors. They were then subcloned into plant expression vector pMDC32 by LR reaction ^41^. GST-Propep1 was amplified from its expression vector and was cloned into pCR8/GW-TOPO to generate pCR8-GST-Propep1 and then subcloned into pMDC32 by LR reaction. The plant expression vectors were transformed into Agrobacterium strain GV3101 and used for infiltration.

Tobacco (*Nicotiana benthamiana*) plants were grown in walk-in-growth chambers at 22°C with a day length of 16 hrs. Tobacco leaves from 4-6 weeks old plants were infiltrated with different gene combinations following a published procedure ^42^. After 3 days, infiltrated leaves were damaged by using forceps as described previously (Hander et al., 2019). After 1 hr of damage, leaf samples were collected and flash frozen in liquid nitrogen. Proteins were extracted with a buffer containing 4M urea, 0.1M Tris, pH 6.8, 1.0% β-ME, and 10 mM EDTA. Extracted proteins were separated in 4-20% precast gradient PAGE gels (Genscript) and transferred to PVDF membranes for immunoblot. The membrane was incubated with GST-TAG antibody (1:1000, ThermoFisher) or anti-MC4 (rabbit, 1:15,000) ^10^. Immunoblots were detected using HRP-conjugated secondary antibodies (for *At*MC4) and SuperSignal West Femto maximum sensitivity substrate (ThermoFisher).

### Mass spectrometry

Wild-type *At*MC4 crystals were harvested in stabilization buffer containing 100 mM sodium cacodylate, pH 6.4, 2.1 M ammonium sulfate. After washing five times with the stabilization buffer, crystals were dissolved in a cleavage reaction solution containing 25 mM HEPES, pH 7.6, 250 mM NaCl, 0.2 mM CaCl_2_ for 10 min at room temperature. The cleaved *At*MC4 fragments were further treated using sequencing-grade chymotrypsin (Roche, Inc.) following manufacturer’s manual. Different from *At*MC4 that cleaves itself at a K or R position, chymotrypsin cleaves at an F, Y, or W position. Digested *At*MC4 peptides was used for mass spectrometry analysis using a Thermo QE-HF (ThermoFisher) at the Stony Brook University Biological Mass Spectrometry Shared Resource. Data was processed using program Proteome Discoverer (ThermoFisher). The cleaved peptides and their positions are listed in Extended Data Table 2.

---

**Extended Data Fig. 1.**
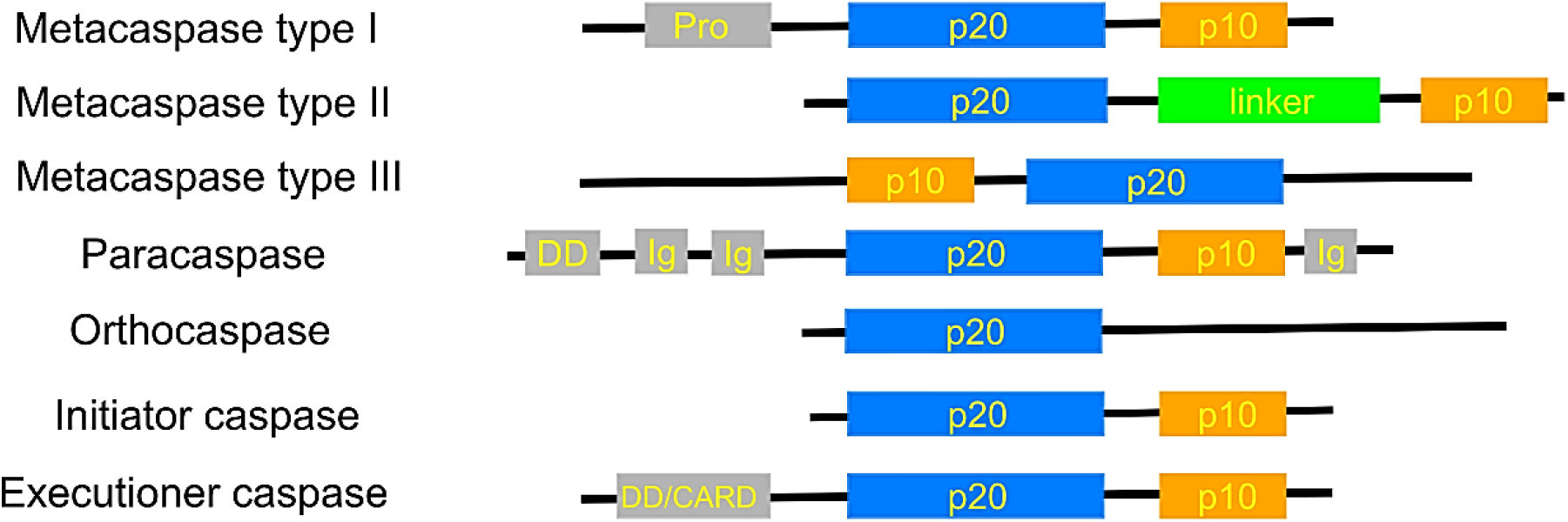
Domain organization of clan CD family 14 cysteine proteases. Caspase core contains the p20 and p10 domains except orthocaspase which has a long C-terminal uncharacterized region. *At*MC4 is a type II metacaspase with a unique linker domain inserted between the p20 and p10 domains.

**Extended Data Fig. 2.**
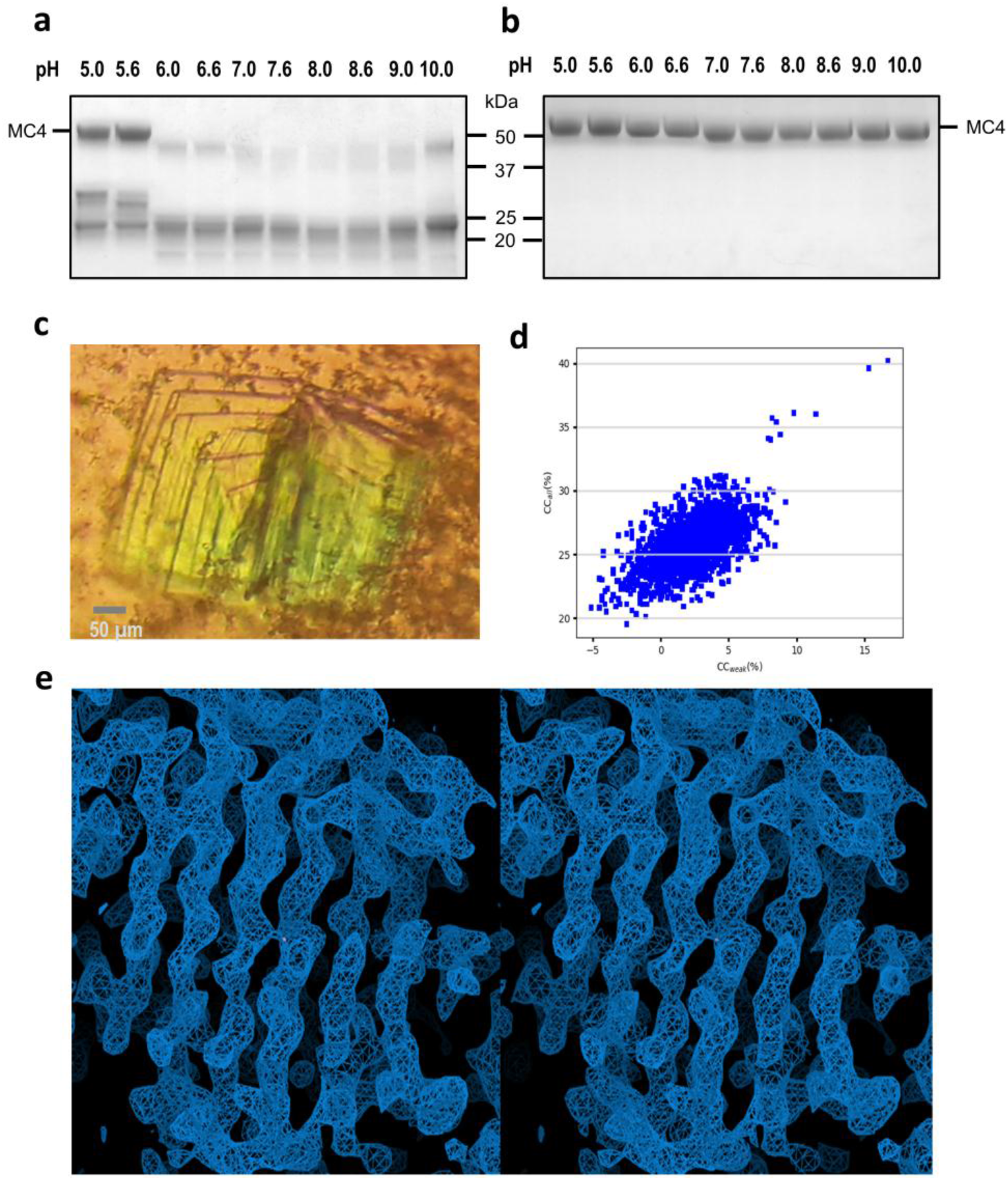
Structure determination of AtMC4 C139A mutant. **a**, pH-dependent cleavage of wild-type *At*MC4 in the presence of 10 mM Ca^2+^. **b**, The catalytic mutant C139A is essentially inactive even in the presence of 10 mM Ca^2+^. **c,** Crystals of the C139A mutant. These crystals formed thin plates (5-10 μm in thickness) and assembled together resembling the pages of an open book. **d**, SHELXD CCweak/CCall plot shows SeMet substructure determination. **e**, A stereo view of the experimental electron densities contoured at 1.0σ for the β-sheet structure in the caspase-like core.

**Extended Data Fig. 3.**
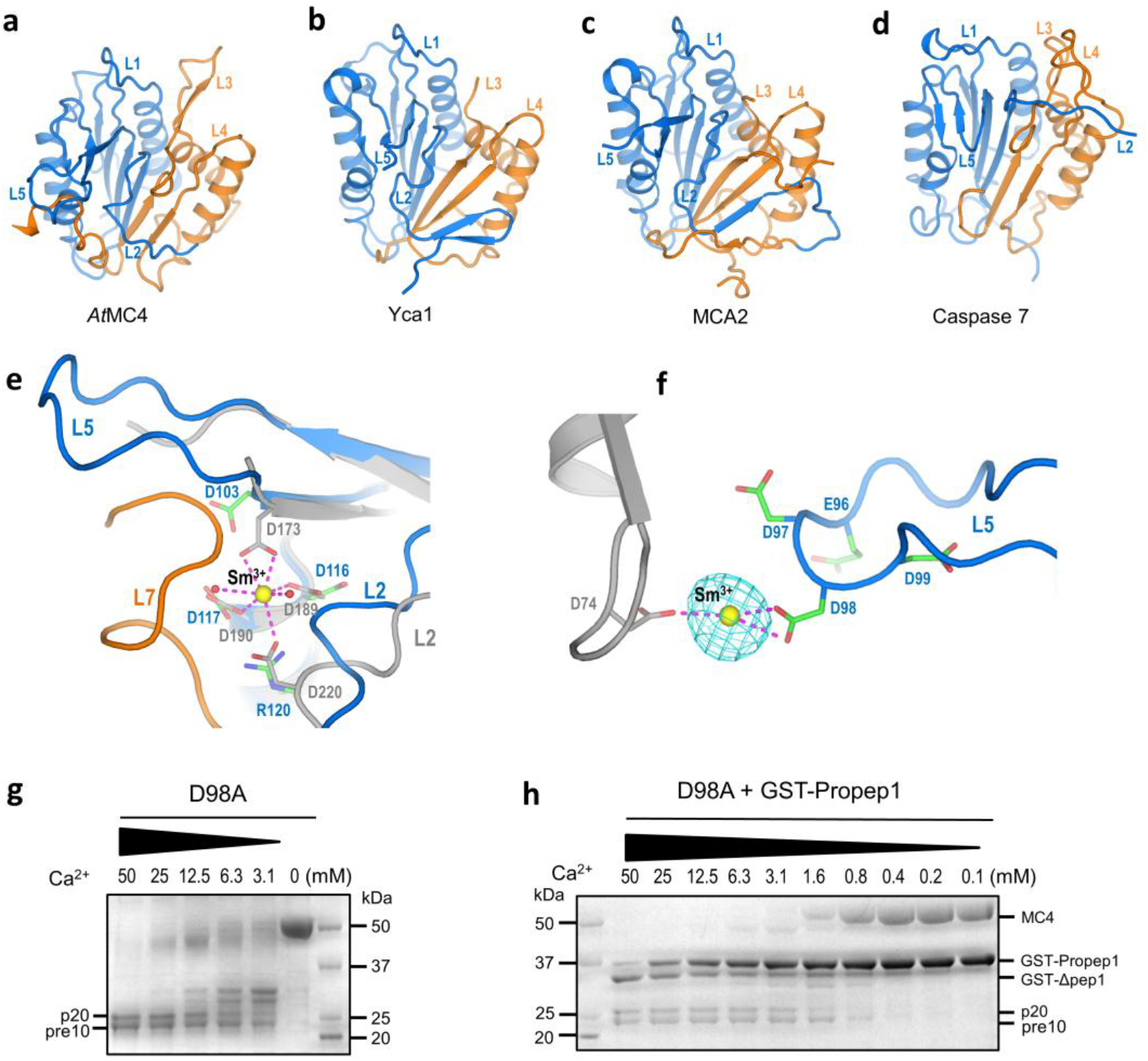
Comparison of the *At*MC4 caspase-like core with its structural relatives and characterization of a Sm^3+^-binding site in *At*MC4. **a**, *At*MC4. **b**, *Saccharomyces cerevisiae* Yca1 (PDB code: 4F6O). **c**, *Trypanosoma brucei* MCA2 (PDB code: 4AFP). **d**, Human Caspase 7 (PDB code: 1K86). The color scheme is marine for p20 and orange for p10 domain. Loops L1 to L5 are indicated. **e**, Structural superimposition of *Trypanosoma brucei* MCA2 (gray) and *At*MC4 for the Sm^3+^-binding site in MCA2. Two water molecules are shown as red spheres. **f**, A different Sm^3+^-binding site in *At*MC4 formed by residue Asp98 in loop L5 and Asp74 from a symmetry-related molecule. The Bijvoet difference Fourier peak for Sm^3+^ is shown as isomeshes contoured at 6σ. **g**, Ca^2+^-dependent self-cleavage in D98A mutant. **h**, Ca^2+^-dependent cleavage of GST-Propep1 by the D98A mutant.

**Extended Data Fig. 4.**
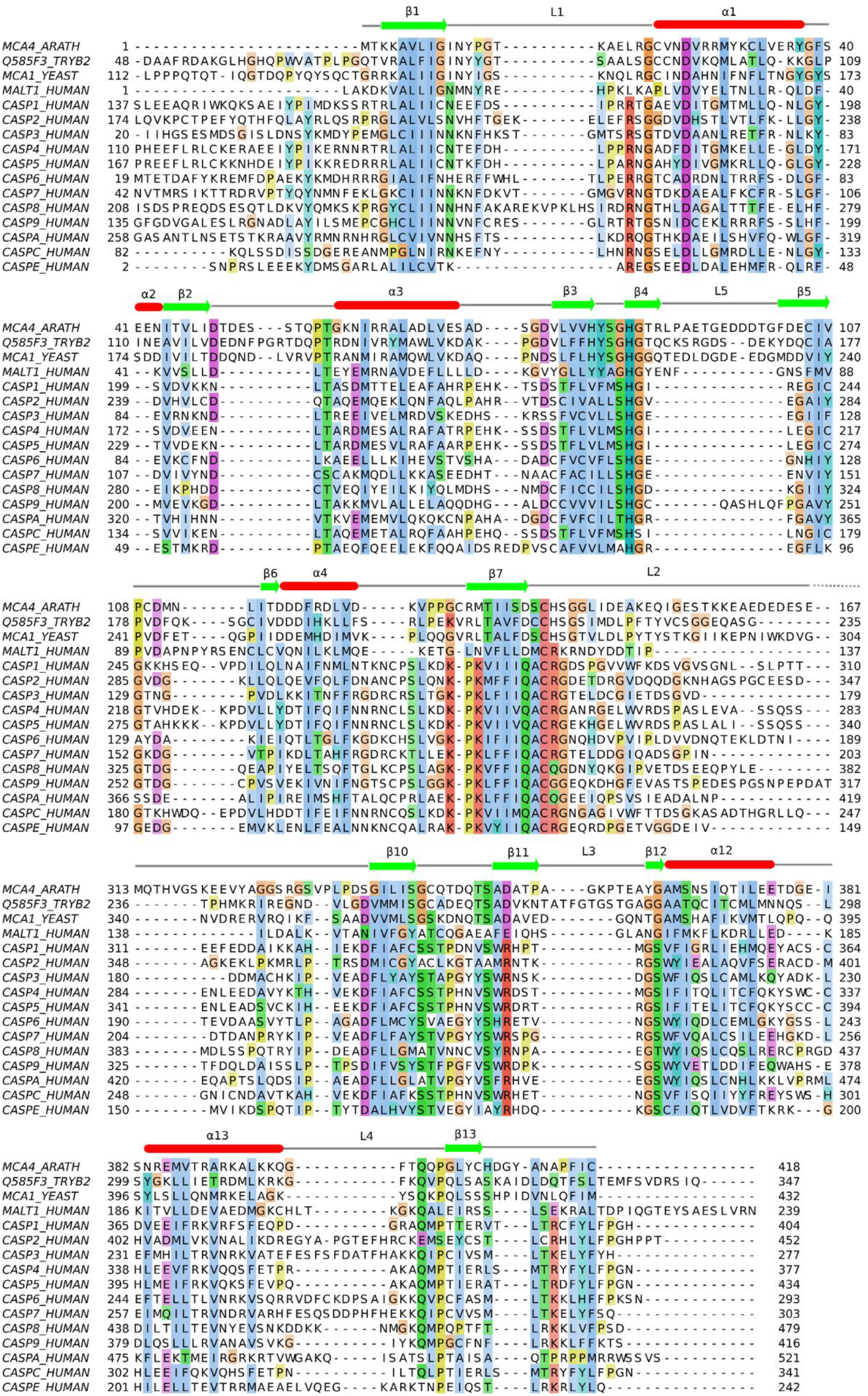
Structure-based sequence alignment of *At*MC4 with its relatives. For simplicity, the linker domain, which is unique to *At*MC4 and type II metacaspases, is not shown in the alignment. In the alignment are UNIPROT (www.uniprot.org) entry names: MCA4_ARATH, *Arabidopsis thaliana MC4; Q585F3_TRYB2: Trypanosoma brucei* MCA2; MCA1_YEAST, Saccharomyces cerevisiae Yca1; MALT1_HUMAN, human MALT1; CASP1_HUMAN, human Caspase 1; CASP2_HUMAN, human Caspase 2; CASP3_HUMAN, human Caspase 3; CASP4_HUMAN, human Caspase 4; CASP5_HUMAN, human Caspase 5; CASP6_HUMAN, human Caspase 6; CASP7_HUMAN, human Caspase 7; CASP8_HUMAN, human Caspase 8; CASP9_HUMAN, human Caspase 9; CASPA_HUMAN, human Caspase 10; CASPC_HUMAN, human Caspase 12; CASPE_HUMAN, human Caspase 14.

**Extended Data Fig. 5.**
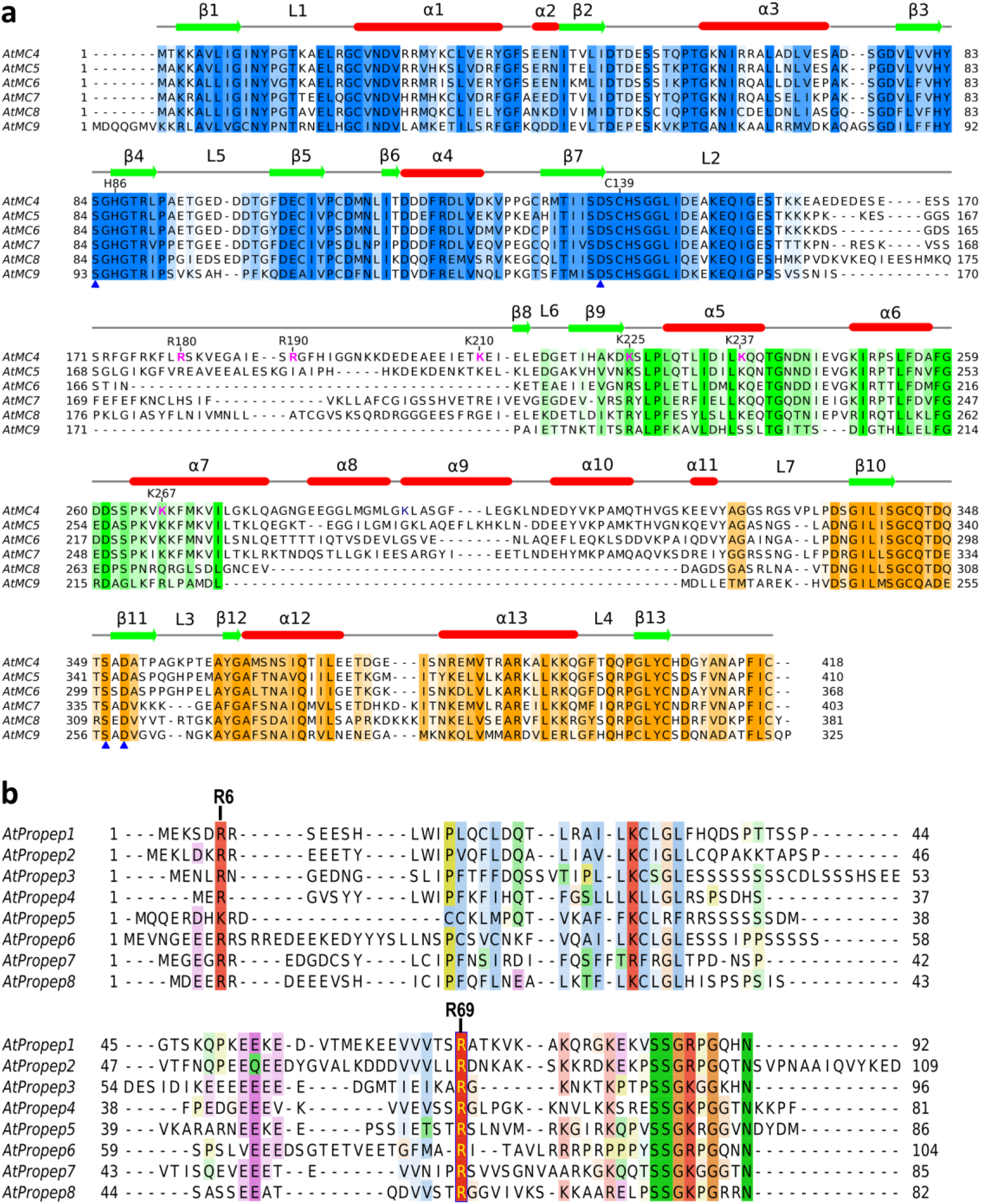
Sequence alignments for type II metacaspases and precursors of elicitor peptide (Propeps) in *Arabidopsis thaliana*. **a,** Structure-based sequence alignment of type II metacaspases in *Arabidopsis thaliana*. Domain coloring is: p20, marine; linker, green; p10, orange. Catalytic residues Cys139 and His86 are indicated. Self-cleavage sites in the linker domain of *At*MC4 are highlighted in magenta. Conserved residues forming the active site are indicated by blue triangles. Residue numbering is for *At*MC4. **b,** Sequence alignment for eight precursors of elicitor peptide in *Arabidopsis thaliana.* Most of these peptides can be processed by *At*MC4 at a conserved position Arg69 ^4^ (*At*Propep1 numbering) in a Ca^2+^-dependent manner. At a higher Ca^2+^ concentration, an additional site at the N-terminus, likely the conserved Arg6 (*At*Propep1 numbering), may be cleaved by *At*MC4.

**Extended Data Fig. 6.**
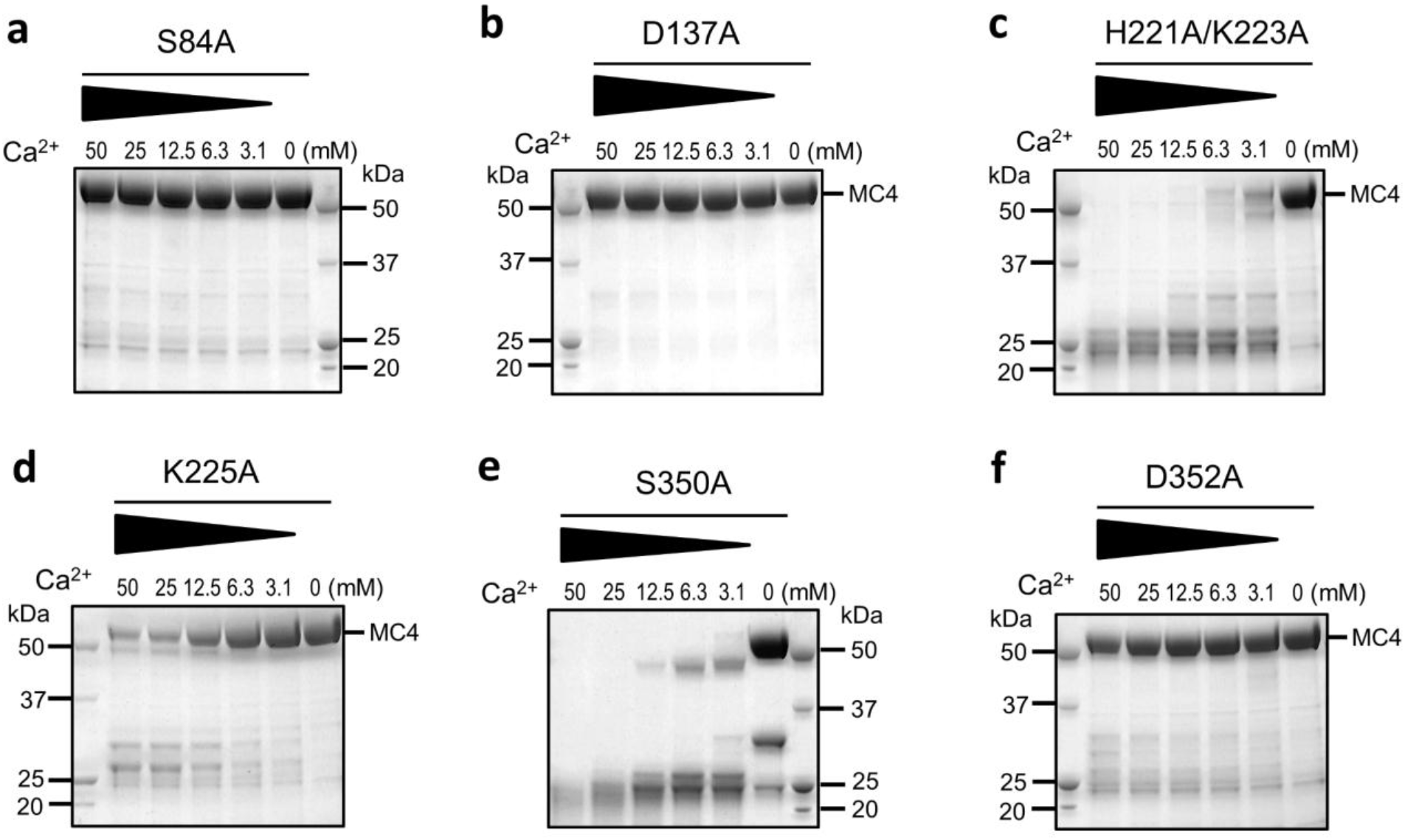
Ca^2+^-dependent self-cleavage for indicated active-site mutants. Indicated mutant proteins were treated with 0-50 mM Ca^2+^ for 10 min followed by SDS-PAGE analysis. Conserved active-site residues S84 (**a**), D137 (**b**), K225 (**d**), and D352 (**f**) are essential for the catalytic activity. Mutating any of them to an alanine essentially abolished self-cleavage activity. Resides H221, K223 (**c**), and S350 (**e**) are on the surface of the active site; and their mutants remain active in self-cleavage.

**Extended Data Fig. 7.**
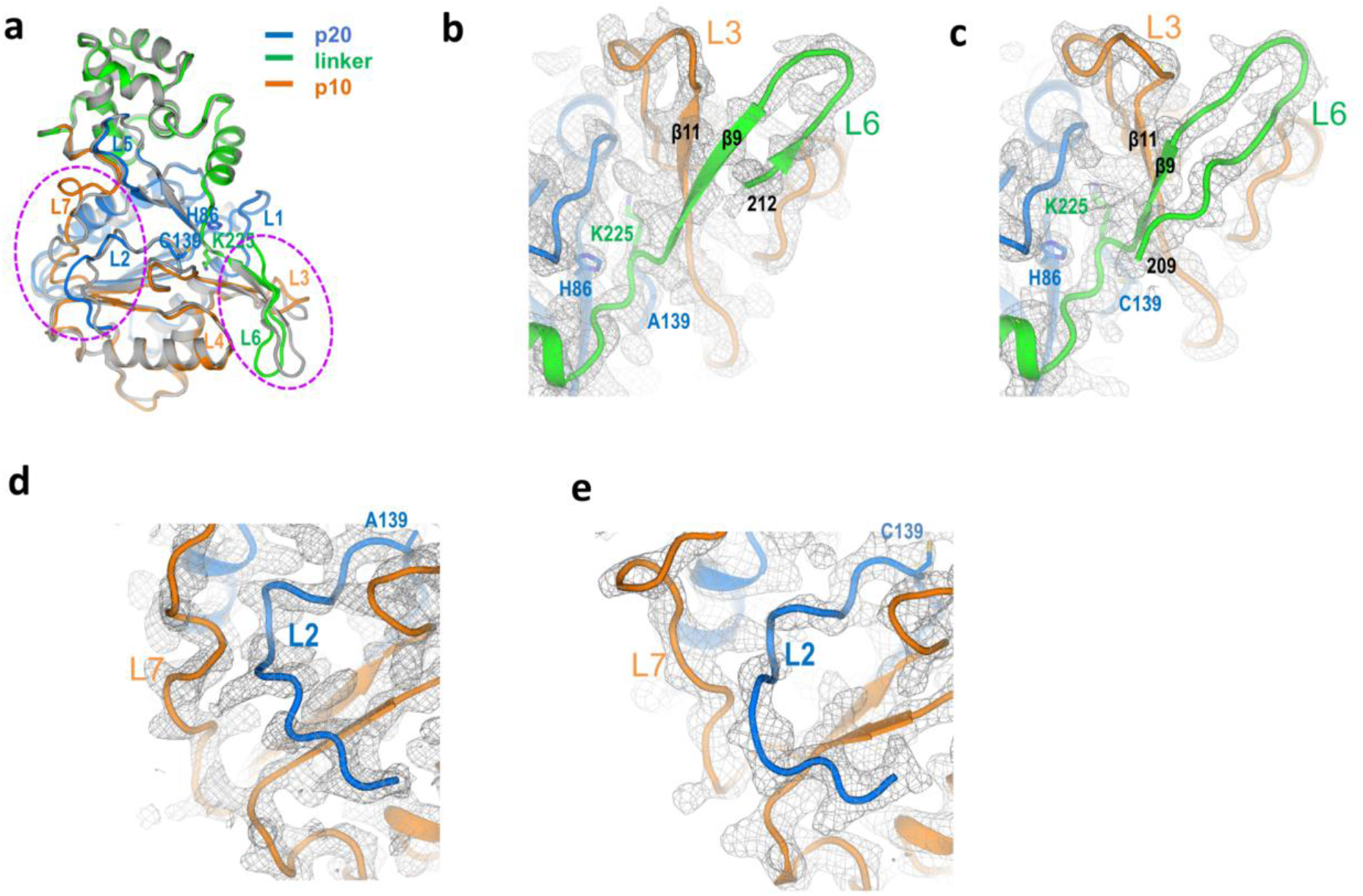
Comparison of wild-type *At*MC4 structure with its C139A mutant, both without Ca^2+^ treatment. **a**, Superimposition of the wild-type and C139A structures. The wild-type structure is shown as cartoons and colored as marine for p20, green for the linker, and orange for p10. The C139A structure is shown as gray cartoons. Dashed ovals indicate the two regions of significant conformational changes between the two structures. The overall structure of the Ca^2+^-free wild-type *At*MC4 is similar to the C139A mutant with an R.M.S.D. of 0.79 Å for 347 aligned Cα atoms. However, significant conformational changes were observed for two regions. One region is loop L6 at the N-terminus of the linker domain and the L3 loop; and the other region is loop L7 at N-terminus of the p10 domain and the L2 loop. These conformational changes might reflect the divergent flexibility in the wild-type and C139A. However, we could not rule out the possibility of crystal packing effects in these observed conformational changes. **b-e**, Electron densities for the flexible loops L3/L6 and L2/L7 in the two structures. In the wild-type structure, ordered electron densities were observed for the region 209-211 that is disordered in the C139A structure. **b**, L3/L6 region in C139A structure. **c**, L3/L6 region in wild-type structure. **d**, L2/L7 region in C139A structure. **e**, L2/L7 region in wild-type structure.

**Extended Data Fig. 8.**
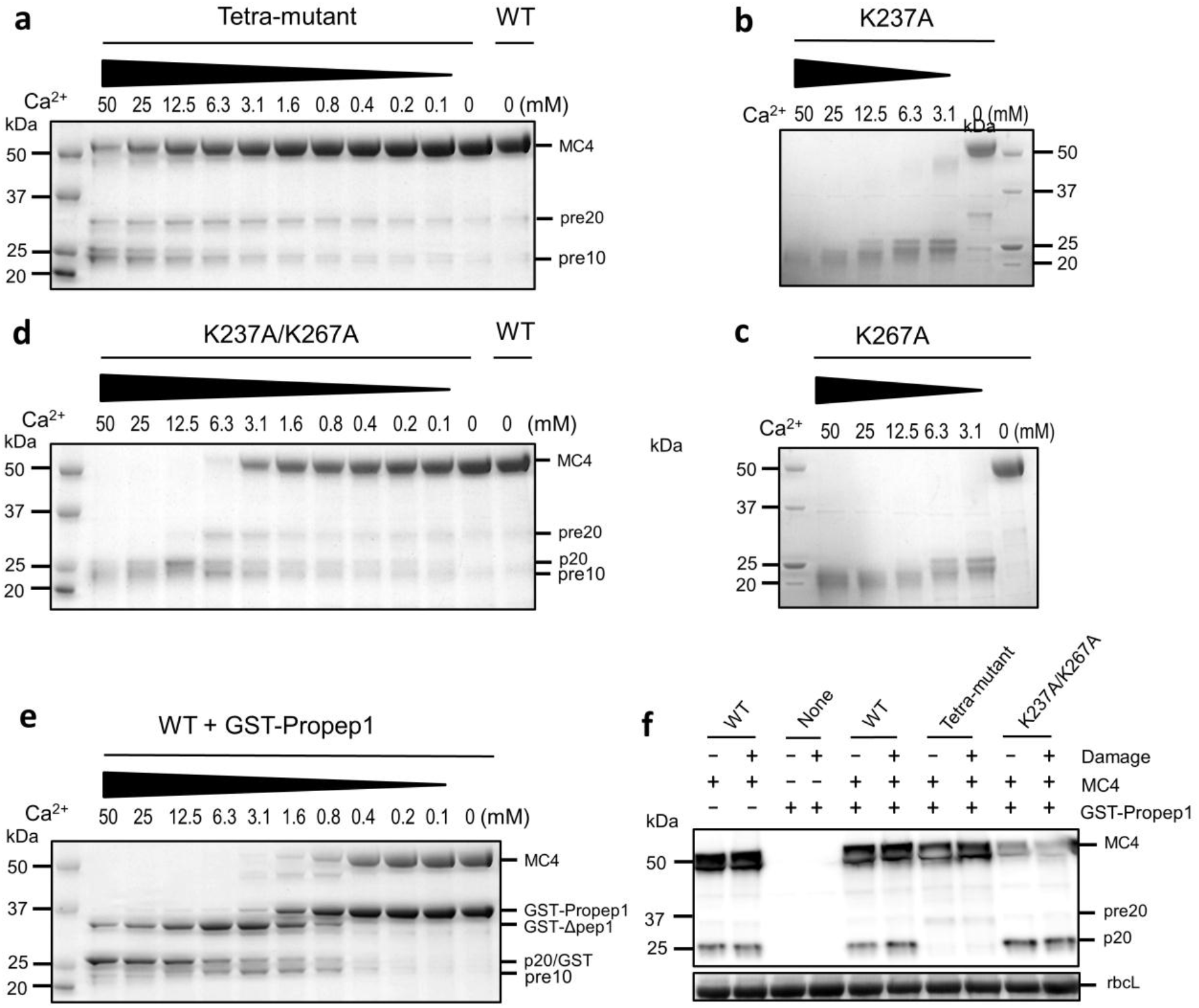
Functional characterization of *At*MC4 activation and substrate processing. **a-d**, Ca^2+^-dependent self-cleavage in *At*MC4 mutants. **a**, 96EDDD99 tetra-mutant. **b**, K237A. **c**, K267A. **d**, K237A/K267A double mutant. **e**, Ca^2+^-dependent activation of *At*MC4 and its cleavage of GST-Propep1. GST-Propep1 was incubated with wild-type *At*MC4 in the presence of Ca^2+^ concentrations ranging from 0 to 50 mM. At low Ca^2+^ concentrations, wild-type *At*MC4 can effectively process GST-Propep1 to release Pep1 peptides. At high Ca^2+^ concentrations (12.5 mM or higher), GST-Δpep1 was further processed at R6/R7 to produce GST with a C-terminal extension. **f**, Damage-induced self-cleavage of *At*MC4 in tobacco (*Nicotiana benthamiana*) leaves. As indicated on top of the lanes, expression vectors containing wild-type *At*MC4 (WT), its various mutants, and GST-Propep1 (+ or −) were transiently expressed in tobacco leaves. Wild-type *At*MC4 and the K237A/K267A double mutant were efficiently cleaved. In contrast, cleavage of the tetra-mutant was mostly suppressed. Data is obtained by immunoblot with a polyclonal antiserum raised against *At*MC4 as described previously (Watanabe and Lam, 2011b). rbcL (the Large subunit of Ribulose 1,5-bisphosphate carboxylase) indicates protein load by Ponceau Red staining.

**Extended Data Table 1.**
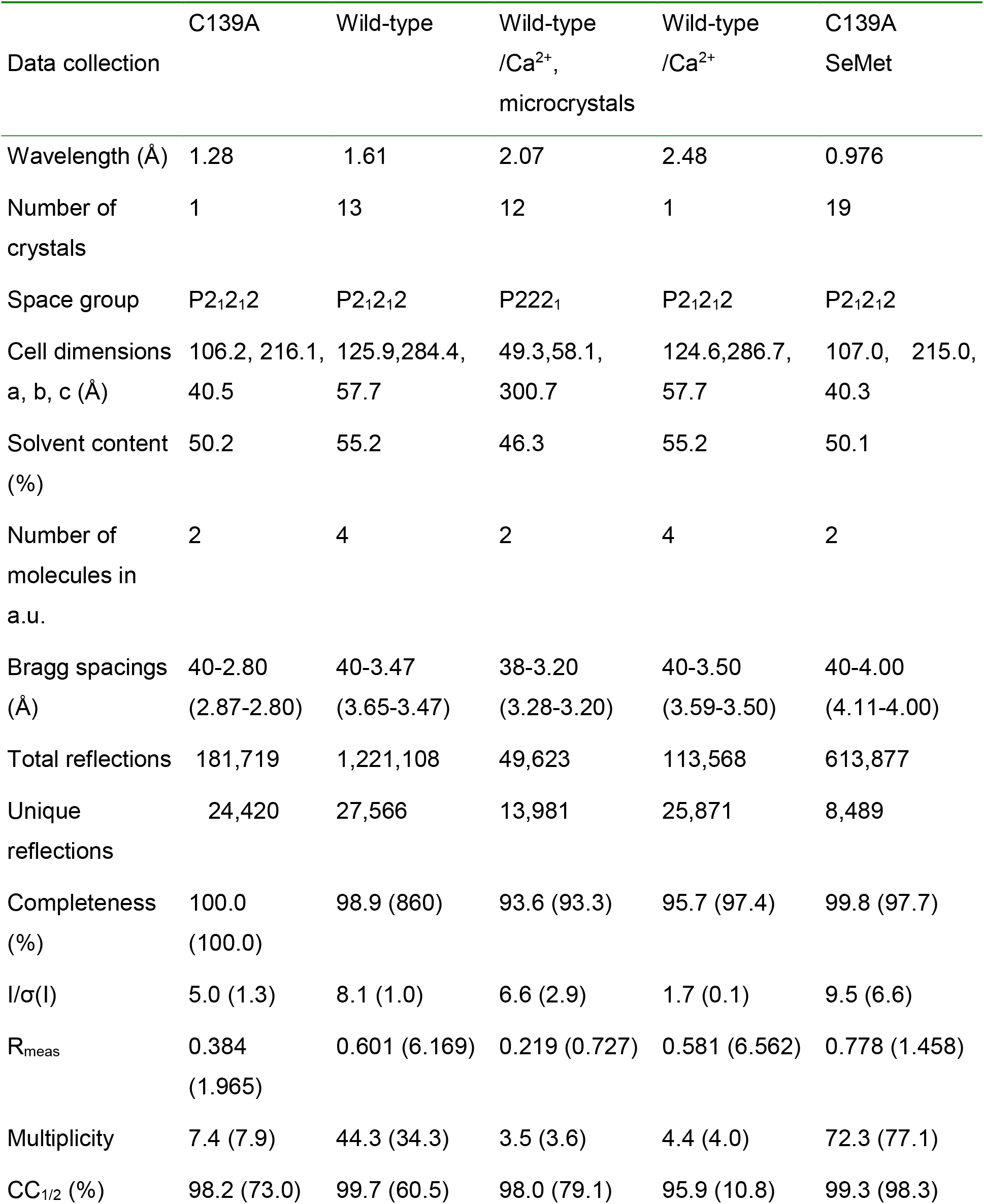
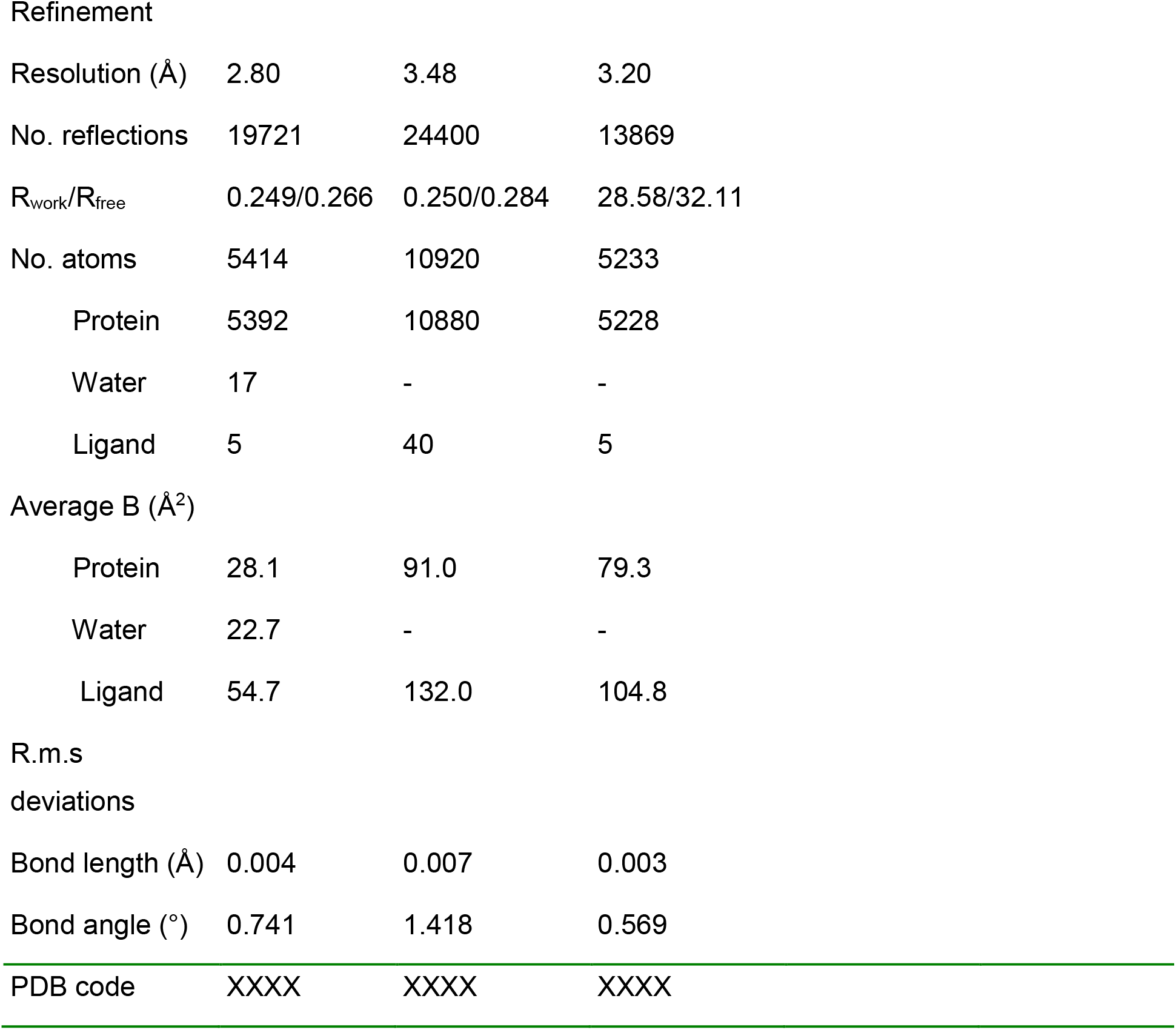
Data collection and refinement statistics.

|  | C139A | Wild-type | Wild-type<br>/Ca<sup>2+</sup>,<br>microcrystals | Wild-type<br>/Ca<sup>2+</sup> | C139A<br>SeMet |
| --- | --- | --- | --- | --- | --- |
| Data collection |  |  |  |  |  |
| Wavelength (Å) | 1.28 | 1.61 | 2.07 | 2.48 | 0.976 |
| Number of<br>crystals | 1 | 13 | 12 | 1 | 19 |
| Space group | P2<sub>1</sub>2<sub>1</sub>2 | P2<sub>1</sub>2<sub>1</sub>2 | P222<sub>1</sub> | P2<sub>1</sub>2<sub>1</sub>2 | P2<sub>1</sub>2<sub>1</sub>2 |
| Cell dimensions<br>a, b, c (Å) | 106.2, 216.1,<br>40.5 | 125.9,284.4,<br>57.7 | 49.3,58.1,<br>300.7 | 124.6,286.7,<br>57.7 | 107.0, 215.0,<br>40.3 |
| Solvent content<br>(%) | 50.2 | 55.2 | 46.3 | 55.2 | 50.1 |
| Number of<br>molecules in<br>a.u. | 2 | 4 | 2 | 4 | 2 |
| Bragg spacings<br>(Å) | 40-2.80<br>(2.87-2.80) | 40-3.47<br>(3.65-3.47) | 38-3.20<br>(3.28-3.20) | 40-3.50<br>(3.59-3.50) | 40-4.00<br>(4.11-4.00) |
| Total reflections | 181,719 | 1,221,108 | 49,623 | 113,568 | 613,877 |
| Unique<br>reflections | 24,420 | 27,566 | 13,981 | 25,871 | 8,489 |
| Completeness<br>(%) | 100.0<br>(100.0) | 98.9 (860) | 93.6 (93.3) | 95.7 (97.4) | 99.8 (97.7) |
| I/σ(I) | 5.0 (1.3) | 8.1 (1.0) | 6.6 (2.9) | 1.7 (0.1) | 9.5 (6.6) |
| R<sub>meas</sub> | 0.384<br>(1.965) | 0.601 (6.169) | 0.219 (0.727) | 0.581 (6.562) | 0.778 (1.458) |
| Multiplicity | 7.4 (7.9) | 44.3 (34.3) | 3.5 (3.6) | 4.4 (4.0) | 72.3 (77.1) |
| CC<sub>1/2</sub> (%) | 98.2 (73.0) | 99.7 (60.5) | 98.0 (79.1) | 95.9 (10.8) | 99.3 (98.3) |

Refinement
|  |  |  |  |
| --- | --- | --- | --- |
| Resolution (Å) | 2.80 | 3.48 | 3.20 |
| No. reflections | 19721 | 24400 | 13869 |
| R<sub>work</sub>/R<sub>free</sub> | 0.249/0.266 | 0.250/0.284 | 28.58/32.11 |
| No. atoms | 5414 | 10920 | 5233 |
| Protein | 5392 | 10880 | 5228 |
| Water | 17 | - | - |
| Ligand | 5 | 40 | 5 |

Average B (Å<sup>2</sup>)
|  |  |  |  |
| --- | --- | --- | --- |
| Protein | 28.1 | 91.0 | 79.3 |
| Water | 22.7 | - | - |
| Ligand | 54.7 | 132.0 | 104.8 |

deviations
|  |  |  |  |
| --- | --- | --- | --- |
| Bond length (Å) | 0.004 | 0.007 | 0.003 |
| Bond angle (°) | 0.741 | 1.418 | 0.569 |

|  |  |  |  |
| --- | --- | --- | --- |
| PDB code | XXXX | XXXX | XXXX |

**Extended Data Table 2.**
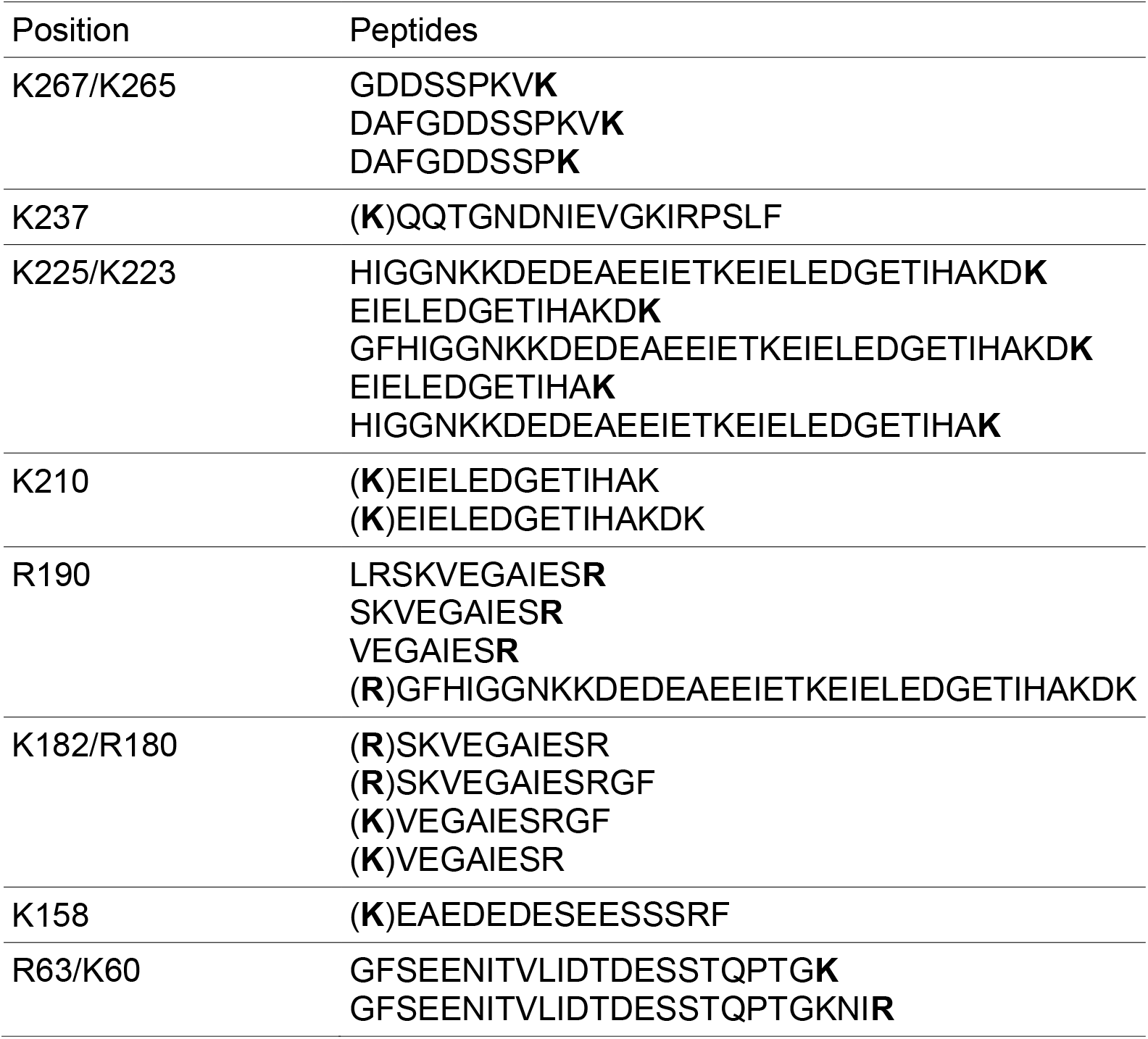
Peptides and their corresponding cleavage site positions identified by mass spectrometry. Self-cleavage sites (R/K) in *At*MC4 are highlighted in bold. R and K in parentheses are self-cleavage sites not present in the sequenced peptides.

## References

1 Suarez, M. F. et al. Metacaspase-dependent programmed cell death is essential for plant embryogenesis. Curr. Biol. 14, R339–R340. (2004).

2 Sundström, J. F. et al. Tudor staphylococcal nuclease is an evolutionarily conserved component of the programmed cell death degradome. Nat. Cell Biol. 11, 1347–1354. (2009).

3 Watanabe, N. & Lam, E. Arabidopsis metacaspase 2d is a positive mediator of cell death induced during biotic and abiotic stresses. Plant J. 66, 969–982. (2011).

4 Hander, T. et al. Damage on plants activates Ca^2+^-dependent metacaspases for release of immunomodulatory peptides. Science 363, eaar7486. (2019).

5 Coll, N. S. et al. Arabidopsis Type I Metacaspases Control Cell Death. Science 330, 1393–1397. (2010).

6 Wen, S. et al. Biochemical evidence of key residues for the activation and autoprocessing of tomato type II metacaspase. FEBS Lett. 587, 2517–2522. (2013).

7 Wong, A. H.-H., Yan, C. & Shi, Y. Crystal Structure of the Yeast Metacaspase Yca1. J. Biol. Chem. 287, 29251–29259. (2012).

8 McLuskey, K. et al. Crystal structure of a Trypanosoma brucei metacaspase. Proc. Natl. Acad. Sci. USA 109, 7469–7474. (2012).

9 Vercammen, D. et al. Type II Metacaspases Atmc4 and Atmc9 of Arabidopsis thaliana Cleave Substrates after Arginine and Lysine. J. Biol. Chem. 279, 45329–45336. (2004).

10 Watanabe, N. & Lam, E. Calcium-dependent activation and autolysis of Arabidopsis metacaspase 2d. J. Biol. Chem. 286, 10027–10040. (2011).

11 Minina, E., Coll, N., Tuominen, H. & Bozhkov, P. Metacaspases versus caspases in development and cell fate regulation. Cell Death Differ. 24, 1314. (2017).

12 Jorgensen, I., Rayamajhi, M. & Miao, E. A. Programmed cell death as a defence against infection. Nat. Rev. Immunol. 17, 151–164. (2017).

13 Uren, A. G. et al. Identification of Paracaspases and Metacaspases: Two Ancient Families of Caspase-like Proteins, One of which Plays a Key Role in MALT Lymphoma. Mol. Cell 6, 961–967. (2000).

14 Klemenčič, M. & Funk, C. Evolution and structural diversity of metacaspases. J. Exp. Bot. 70, 2039–2047. (2019).

15 Shi, Y. Caspase activation: revisiting the induced proximity model. Cell 117, 855–858. (2004).

16 Clapham, D. E. Calcium signaling. Cell 80, 259–268. (1995).

17 Knight, H. in Int. Rev. Cytol. Vol. 195 269–324 (Elsevier, 1999).

18 Xiong, L., Schumaker, K. S. & Zhu, J.-K. Cell signaling during cold, drought, and salt stress. Plant Cell 14, S165–S183. (2002).

19 Zhang, L., Du, L. & Poovaiah, B. Calcium signaling and biotic defense responses in plants. Plant Signal. Behav. 9, e973818. (2014).

20 McAinsh, M. R. & Pittman, J. K. Shaping the calcium signature. New Phytol. 181, 275–294. (2009).

21 Holm, L. & Rosenstrom, P. Dali server: conservation mapping in 3D. Nucleic Acids Res. 38, W545–W549. (2010).

22 Chai, J. et al. Crystal Structure of a Procaspase-7 Zymogen: Mechanisms of Activation and Substrate Binding. Cell 107, 399–407. (2001).

23 Guo, G. et al. Synchrotron microcrystal native-SAD phasing at a low energy. IUCrJ 6. (2019).

24 Tang, J. et al. Structural basis for recognition of an endogenous peptide by the plant receptor kinase PEPR1. Cell Res. 25, 110–120. (2015).

25 Salvesen, G. S. & Dixit, V. M. Caspase activation: the induced-proximity model. Proc. Natl. Acad. Sci. USA 96, 10964–10967. (1999).

26 Edel, K. H., Marchadier, E., Brownlee, C., Kudla, J. & Hetherington, A. M. The Evolution of Calcium-Based Signalling in Plants. Curr Biol 27, R667–R679. (2017).

27 Jiang, Z. et al. Plant cell-surface GIPC sphingolipids sense salt to trigger Ca(2+) influx. Nature 572, 341–346. (2019).

28 Tian, W. et al. A calmodulin-gated calcium channel links pathogen patterns to plant immunity. Nature 572, 131–135. (2019).

29 Baker, N. A., Sept, D., Joseph, S., Holst, M. J. & McCammon, J. A. Electrostatics of nanosystems: Application to microtubules and the ribosome. P Natl Acad Sci USA 98, 10037–10041. (2001).

30 Jeong, J.-Y. et al. One-step sequence-and ligation-independent cloning as a rapid and versatile cloning method for functional genomics studies. Appl. Environ. Microbiol. 78, 5440–5443. (2012).

31 Winter, G. et al. DIALS: implementation and evaluation of a new integration package. Acta Cryst. D 74, 85–97. (2018).

32 Winn, M. D. et al. Overview of the CCP4 suite and current developments. Acta Cryst. D 67, 235–242. (2011).

33 Schneider, T. R. & Sheldrick, G. M. Substructure solution with SHELXD. Acta Cryst. D 58, 1772–1779. (2002).

34 Vonrhein, C., Blanc, E., Roversi, P. & Bricogne, G. Automated structure solution with autoSHARP. Methods Mol. Biol. 364, 215–230. (2007).

35 Cowtan, K. The Buccaneer software for automated model building. 1. Tracing protein chains. Acta Cryst. D 62, 1002–1011. (2006).

36 Afonine, P. V. et al. Towards automated crystallographic structure refinement with phenix.refine. Acta Cryst. D 68, 352–367. (2012).

37 Emsley, P., Lohkamp, B., Scott, W. G. & Cowtan, K. Features and development of Coot. Acta Cryst. D 66, 486–501. (2010).

38 Laskowski, R. A., Macarthur, M. W., Moss, D. S. & Thornton, J. M. Procheck - a program to check the stereochemical quality of protein structures. J. Appl. Cryst. 26, 283–291. (1993).

39 Chen, V. B. et al. MolProbity: all-atom structure validation for macromolecular crystallography. Acta Cryst. D 66, 12–21. (2010).

40 Liu, Q. et al. Structures from Anomalous Diffraction of Native Biological Macromolecules. Science 336, 1033–1037. (2012).

41 Curtis, M. D. & Grossniklaus, U. A gateway cloning vector set for high-throughput functional analysis of genes in planta. Plant Physiology 133, 462–469. (2003).

42 Ohad, N. & Yalovsky, S. Utilizing bimolecular fluorescence complementation (BiFC) to assay protein-protein interaction in plants. Methods Mol Biol 655, 347–358. (2010).

